# Resolving Challenges in Detection and Quantification of D-2-hydroxyglutarate and L-2-hydroxyglutarate via LC/MS

**DOI:** 10.1101/2024.04.26.591335

**Authors:** Tyrone Dowdy, Mioara Larion

## Abstract

D-2-Hydroxyglutarate and L-2-Hydroxyglutarate (D-2HG/L-2HG) are typically metabolites of non-specific enzymatic reactions that are kept in check by the housekeeping enzymes, D-2HG /L-2HG dehydrogenase (D-2HGDH/L-2HGDH). In certain disease states, such as D-2HG or L-2HG aciduria and cancers, accumulation of these biomarkers interferes with oxoglutarate-dependent enzymes that regulate bioenergetic metabolism, histone methylation, post-translational modification, protein expression and others. D-2HG has a complex role in tumorigenesis that drives metabolomics investigations. Meanwhile, L-2HG is produced by non-specific action of malate dehydrogenase and lactate dehydrogenase under acidic or hypoxic environments. Characterization of divergent effects of D-2HG/L-2HG on the activity of specific enzymes in diseased metabolism depends on their accurate quantification via mass spectrometry. Despite advancements in high-resolution quadrupole time-of-flight mass spectrometry (HR-QTOF-MS), challenges are typically encountered when attempting to resolve of isobaric and isomeric metabolites such as D-2HG/L-2HG for quantitative analysis. Herein, available D-2HG/L-2HG derivatization and liquid chromatography (LC) MS quantification methods were examined. The outcome led to the development of a robust, high-throughput HR-QTOF-LC/MS approach that permits concomitant quantification of the D-2HG and L-2HG enantiomers with the benefit to quantify the dysregulation of other intermediates within interconnecting pathways. Calibration curve was obtained over the linear range of 0.8-104 nmol/mL with r^2^ ≥ 0.995 for each enantiomer. The LC/MS-based assay had an overall precision with intra-day CV % ≤ 8.0 and inter-day CV % ≤ 6.3 across the quality control level for commercial standard and pooled biological samples; relative error % ≤ 2.7 for accuracy; and resolution, R_s_= 1.6 between 2HG enantiomers (m/z 147.030), D-2HG and L-2HG (at retention time of 5.82 min and 4.75 min, respectively) following chiral derivatization with diacetyl-L-tartaric anhydride (DATAN). Our methodology was applied to disease relevant samples to illustrate the implications of proper enantioselective quantification of both D-2HG and L-2HG. The stability of the method allows scaling to large cohorts of clinical samples in the future.

## Introduction

The enhanced ability to resolve the D- and L-enantiomers of 2-hydroxyglutarate (2HG) racemic mixture is vital to clinically differentiate diseases such as D-/L-2-hydroxyglutaric aciduria, the presence of isocitrate dehydrogenase mutations in gliomas, acute myeloid leukemia, chondrosarcomas, cholangiocarcinoma’s, in addition to accurately quantify, elucidate and differentiate their biologically roles and potency [1–5].

D-2HG and L-2HG are produced as normal metabolites from nonspecific enzymatic activities of phosphoglycerate, malate and lactate dehydrogenases particularly under hypoxic conditions (Figure 1) [6,7]. High levels of D-2HG appear as the result of novel catalytic activity of mutant isocitrate dehydrogenase (IDH*^mut^*) 1 and 2 in the above-mentioned tumors, or due to loss of function mutations of D-2hydroxyglutarate dehydrogenase (D-2HGDH) associated with D-2hydroxyglutaric aciduria, a neurometabolic disorder [8,9]. Importantly, elevated D-2HG concentration serves as an oncomarker for the activity of IDH*^mut^* enzymes in tumor development; thus, it is used to diagnose, assess prognosis and determine treatment interventions. High levels of L-2HG arise mainly from loss of function mutations in the housekeeping enzyme, L-2-hydroxyglutarate dehydrogenase (L-2HGDH) and are used to identify L-2HG-induced aciduria, which sometimes leads to gliosis and tumorigenesis via unknown mechanisms [10]. Under typical LC/MS conditions, the co-elution of isomeric compounds D-2HG and L-2HG causes limitations in differentiating their contribution to microenvironments when standard method allow only the total 2HG racemic mixture to be measured [11].

**Figure 1.**
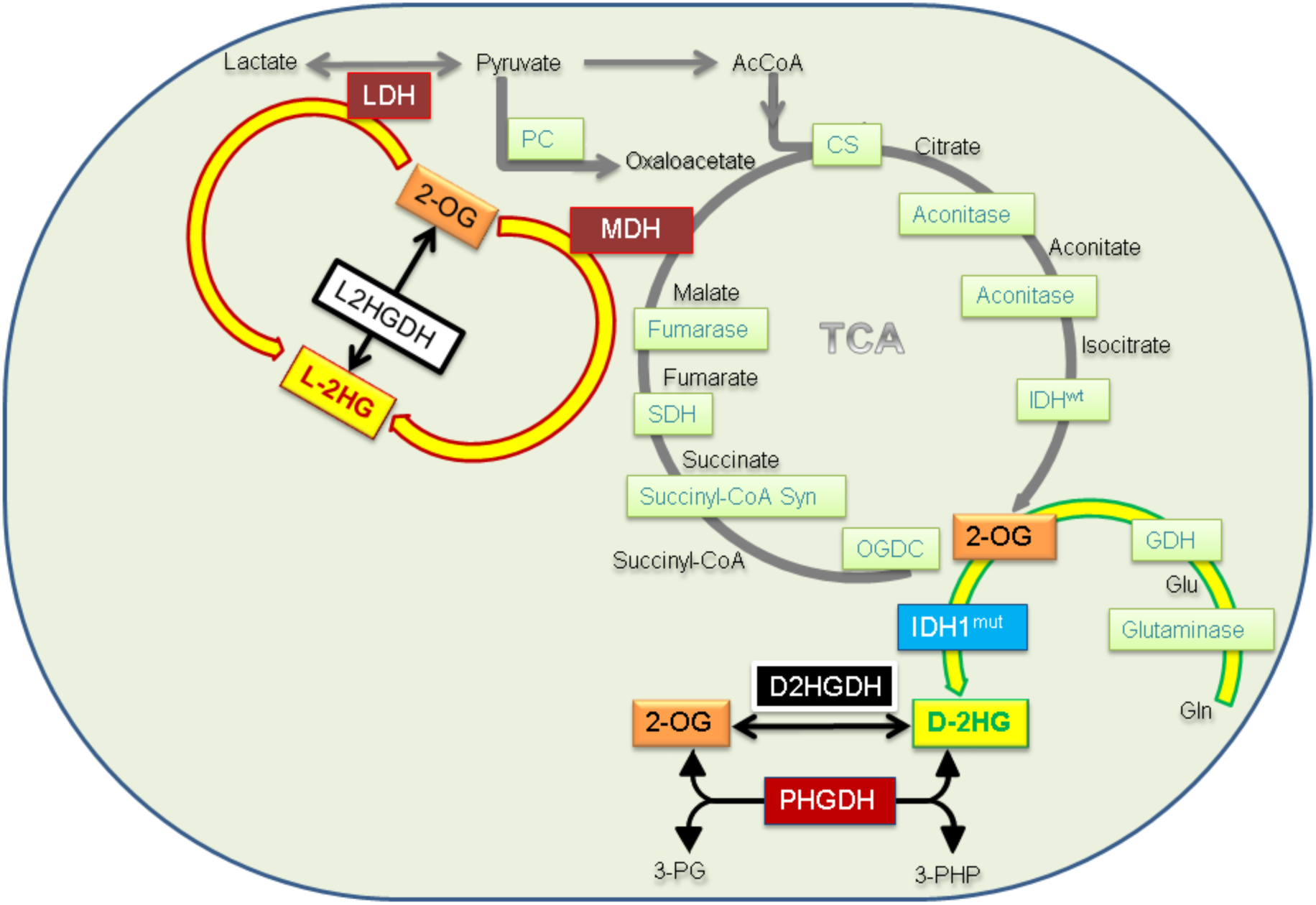
Independent production of D-2HG and L-2HG by normal, mutant and nonspecific enzymatic activity. 2- OG, 2-oxoglutarate; 3-PG, 3-phosphoglycerate; 3-PHP, phosphohydroxypyruvate; AcCoA, Acetyl-Coenzyme A; CS, citrate synthase; D2HGDH, D-2HG dehydrogenase ; GDH, glutamate dehydrogenase; Gln, glutamine; Glu, Glutamate; IDH^wt^, Isocitrate dehydrogenase; IDH1^mut^, mutant IDH1; L2HGDH, L-2HG dehydrogenase; LDH, lactate dehydrogenase; MDH, malate dehydrogenase; OGDC, oxoglutarate dehydrogenase complex; PC, pyruvate carboxylase; PDH, pyruvate dehydrogenase; PHGDH, phosphoglycerate dehydrogenase; SDH, succinate dehydrogenase; Syn, synthase; TCA, tricyclic acid cycle.

The importance of robust D-2HG and L-2HG quantification is linked with both clinical diagnosis of above-mentioned diseases as well as the understanding of underlying mechanisms that lead to diseases induced by these two metabolites, as highlighted in Table 1. For example, in gliomas, there is an ongoing debate regarding the hypothesis that D-2HG either stabilizes HIF1α and promotes Warburg effect, or triggers HIF1α degradation [12–14]. In addition, L-2HG has been reported to stabilize HIF1α [15].

**Table 1.**
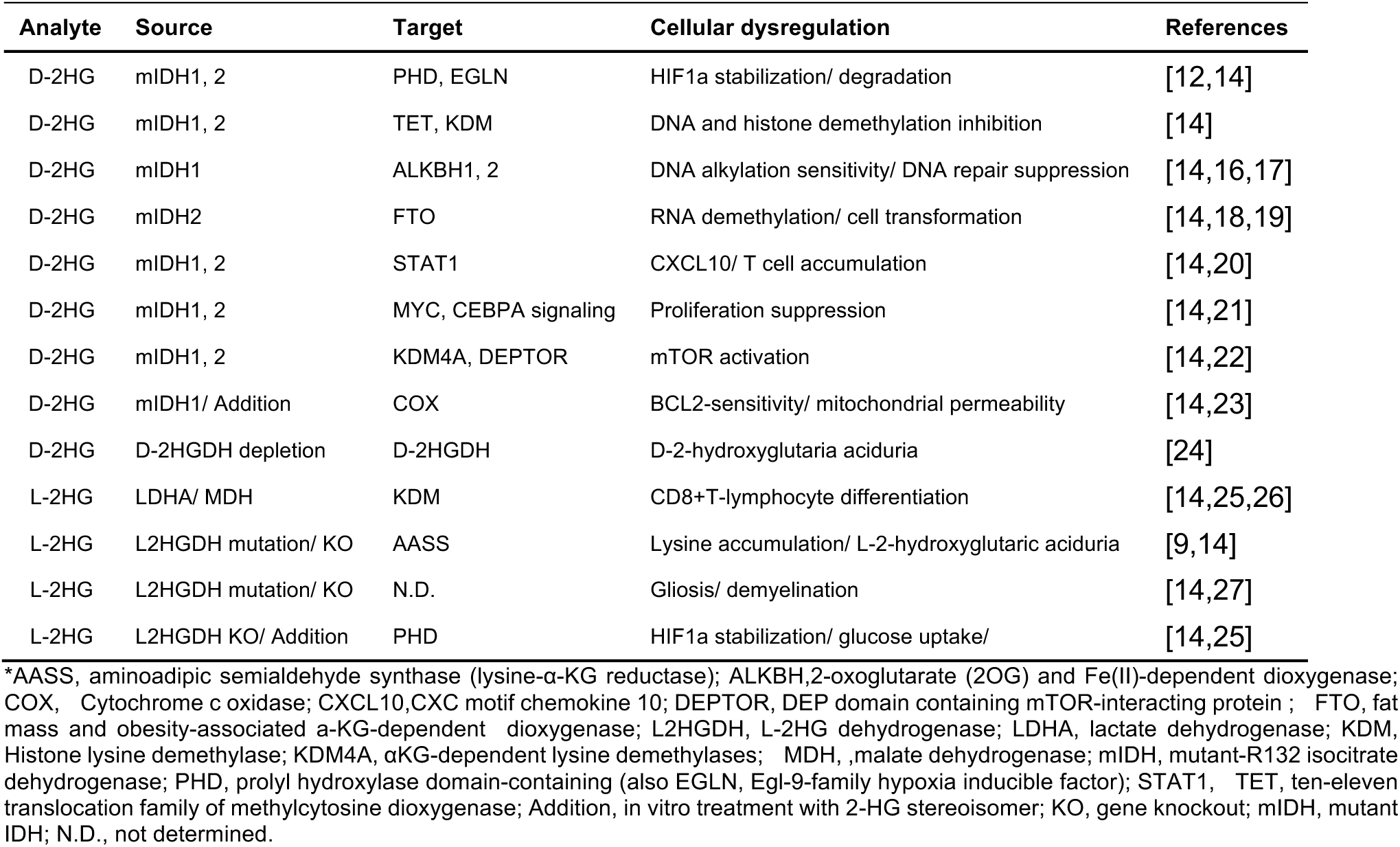
Differential role of 2HG-stereoisomers in cellular regulation*.

Earlier studies suggested that D-2HG and L-2HG, stereoisomers possess similar roles in mediating specific 2-oxoglutarate dependent enzymes at variable degrees of potency [7]. In addition, studies have revealed that certain biological conditions (e.g., low pH, hypoxia, normoxia, genetic mutation and orientation of functional groups) impact the production and activity of specific 2HG enantiomer [7]. The slight variation in structure of enantiomers can profoundly impact the binding affinity, specificity, reactivity (e.g. nucleophilic substitution), and steric interactions for bioactive molecules. These distinctions exemplify the importance of nonspecific enzyme-substrate activity stimulate the selective overproduction of D-2HG or L-2HG enantiomers (Table 1). Although these 2HG stereoisomers present identical mass, chemical formula, composition, and 2D-chemical structure, the presence of chiral carbon generates asymmetry which facilitates independent quantification of stereoisomers that would co-elute without chiral derivatization which is necessary to accurately determine concentrations, measure changes in their regulation and differentiate these biological effects.

While previous LC/MS-based studies involving derivatization with N-p-tosyl-L-phenylalanyl chloride (TSPC) have shown selectivity in separation of D-2HG and L-2HG, our efforts to reproduce revealed challenges with the sample stability and presence of variable amount of underivatized D-2HG and L-2HG. Given that 2HG enantiomers tend to co-elute under nonchiral conditions (in underivatized samples injected on a nonchiral column), this prompted us to re-examine the existing derivatization and LC/MS methods involving the use of DATAN derivatizing agent in order to make them robust enough for large scale clinical applications [28,29]. Herein, we report an optimized methodology for accurate, simultaneous quantification of both 2HG enantiomers. We combined the stability of diacetyl-L-tartaric anhydride (DATAN)-2HG mixture, with the ability of DATAN to resolve the enantiomers chromatographically and the advantage to measure the D-2HG and L-2HG analogs (m/z 147.030) of the DATAN ester (m/z 363.057). We compared six columns with different chemistry, compound standards from different vendors, derivatization approaches as well as incorporated unique solvent conditions and sample preparation in order to determine an ideal combination for optimal chromatography. This strategy also permitted simultaneous detection of other disease relevant metabolites. Our study confirmed that a strong correlation exists between changes in glutamine levels that directly correspond to overproduction of D-2HG specifically; furthermore, provided evidence in support of the hypothesis that mutant IDH1 gliomas utilize glutamine to supplement TCA cycle.

## Results

### Derivatization Optimization for 2-Hydroxyglutaric Acid Racemic Mixture

Our derivatization method is based on chemical reaction with diacetyl-L-tartaric anhydride (DATAN). We introduced steric constraints to improve resolution for 2HG enantiomers for LC/MS quantification (Figure 2) [30]. The racemic mixture of 2HG (2HG-rac) was subjected to derivatization agents, (e.g. diacetyl-L-tartaric anhydride, DATAN) under acidic conditions while applying heat which activates the condensation reaction shown in Figure 2.

**Figure 2.**
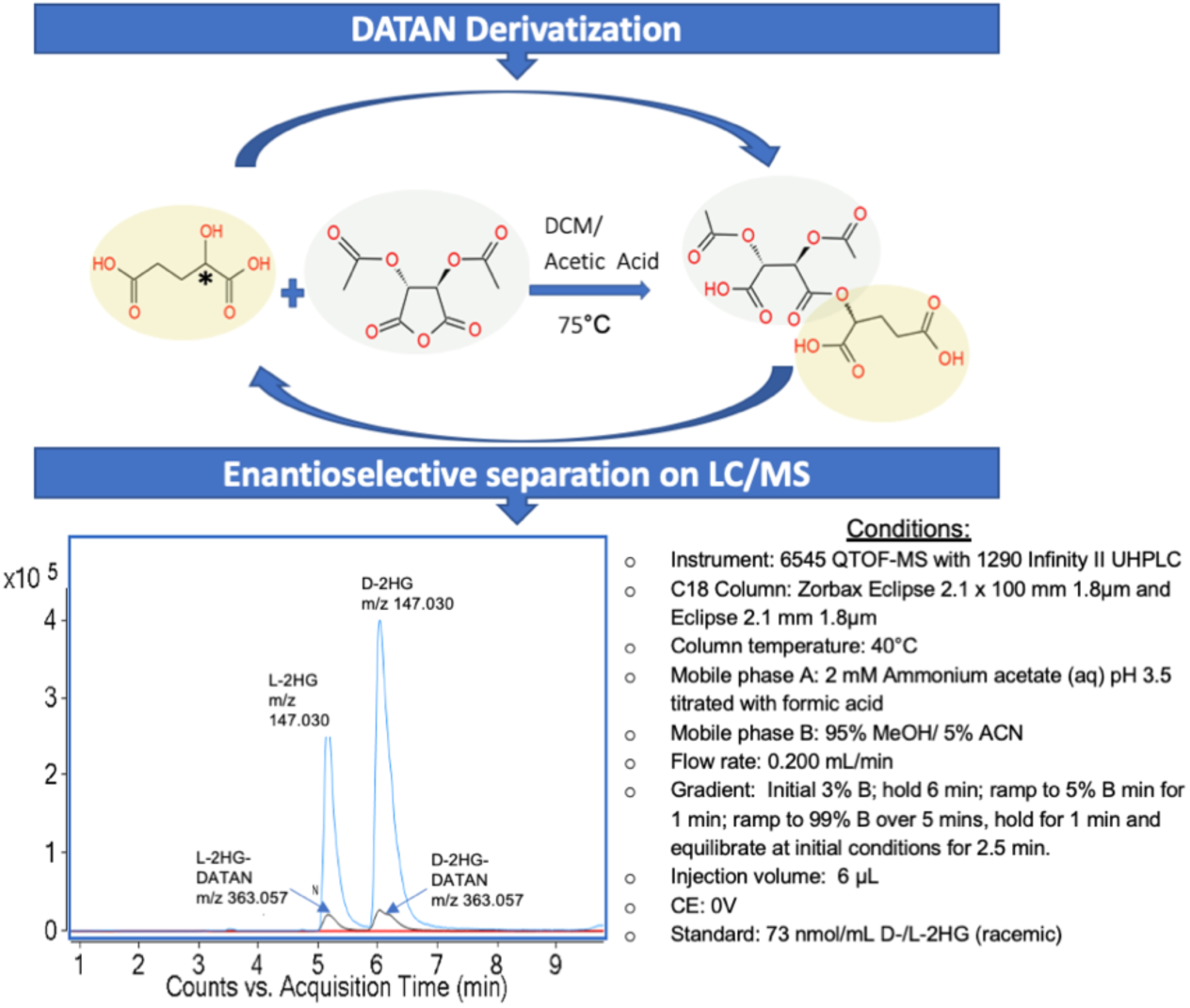
The strategy for D-2HG and L-2HG separation and quantification with DATAN and Agilent 6545 LC/MS QTOF. The derivatization with DATAN facilitates chromatographic separation of D-/L-2HG-DATAN esters (m/z 363.057) with nonchiral C18 column using conditions detailed in the Results section. Following separation, the esters are fragmented during MS analysis which facilitates direct quantification of the D- and L-2HG analogs (m/z 147.030). The racemic standard of D- and L-2HG were derivatized with DATAN, then analyzed via LC/MS as described in Method section. D- and L-2HG standards were injected and their XICs were extracted within ±5 ppm.

The resulting DATAN esterification shown in Figure 2 induces steric strain at the chiral center which likely contributes to an additional dimension of separation and differential elution of enantiomers along nonchiral analytical columns [18]. There are a series of qualitative advantages to conducting derivatization with DATAN. Unlike 2HG-N-p-tosyl-L-phenylalanyl chloride (TSPC) esters, 2HG-DATAN esters present a chromophore that serves as a visible yellow indicator for the esterification reaction [31]. As a result, there is a physical change in appearance of the extract from clear to yellow following derivatization with DATAN. This provides a fast-qualitative assessment of experiment prior to proceeding with LC/MS analysis. Samples that are saturated with organic acids or other reaction sites will appear brown, which can provide a preliminary indication a problematic experiment. In the latter case, it is beneficial to dilute the sample and only derivatize an aliquot to ensure chemical equilibrium with the total 2HG derivatization in the sample. The DATAN ester of 2HG (2HG-DATAN) fragments during MS acquisition which enables the exclusive quantification of the 2HG transition (m/z 147.030) and the 2HG-DATAN precursor (m/z 363.057) (Figure 2).

### Detection of D-2HG and L-2HG under the optimized LC-MS quantification methodology

While TOF-MS (time-of-flight mass spectrometry) detection favors the highly abundant m/z 147.030 (2HG) over m/z 363.057 (2HG-DATAN) when collision induces dissociation (CID) at 0V or greater, it was inferred that m/z 147.030 selection is likely due to in-source fragmentation during MS or MS/MS mode. Given that the ionization of the 2HG moiety exceeds that of 2HG-DATAN ester, quantification was based on m/z 147.030 as the quant ion relying on the high resolution, accurate mass capabilities of quadruple time-of-flight MS without applying tandem MS (MS/MS) (Figure 2). Single quadrupole time-of-flight mass spectrometer (QTOF-MS) experience challenges to achieve the sensitivity, or the ion monitoring scan rates of the triple quadrupole mass spectrometers (QQQ), typically, used for multiple-reaction monitoring MRM [32]. Unlike QQQ which has 3 mass filters, single Q-TOF-MS permits a wider mass spectra extraction window during tandem target MS/MS, which can limit the scanning resources and compromised sensitivity [32,33]. Therefore, an accurate mass TOF MS-based approach was selected on the high-resolution instrument in order to enhance detection and facilitate the incorporation of targeted profiling with validated analytes.

### Detection and separation from endogenous isomers

In the next step, we analyzed the TOF-MS data to determine whether accurate separation and quantification of 2HG moiety from the DATAN ester can be achieved without interfering metabolites. Target spectra were inspected from a derivatized blank (composed of diluent only) and pure standards from each potential endogenous isomer of 2HG baring identical accurate mass, m/z and/or chemical composition (Figure 3 and Supplementary Figure S1).

**Figure 3.**
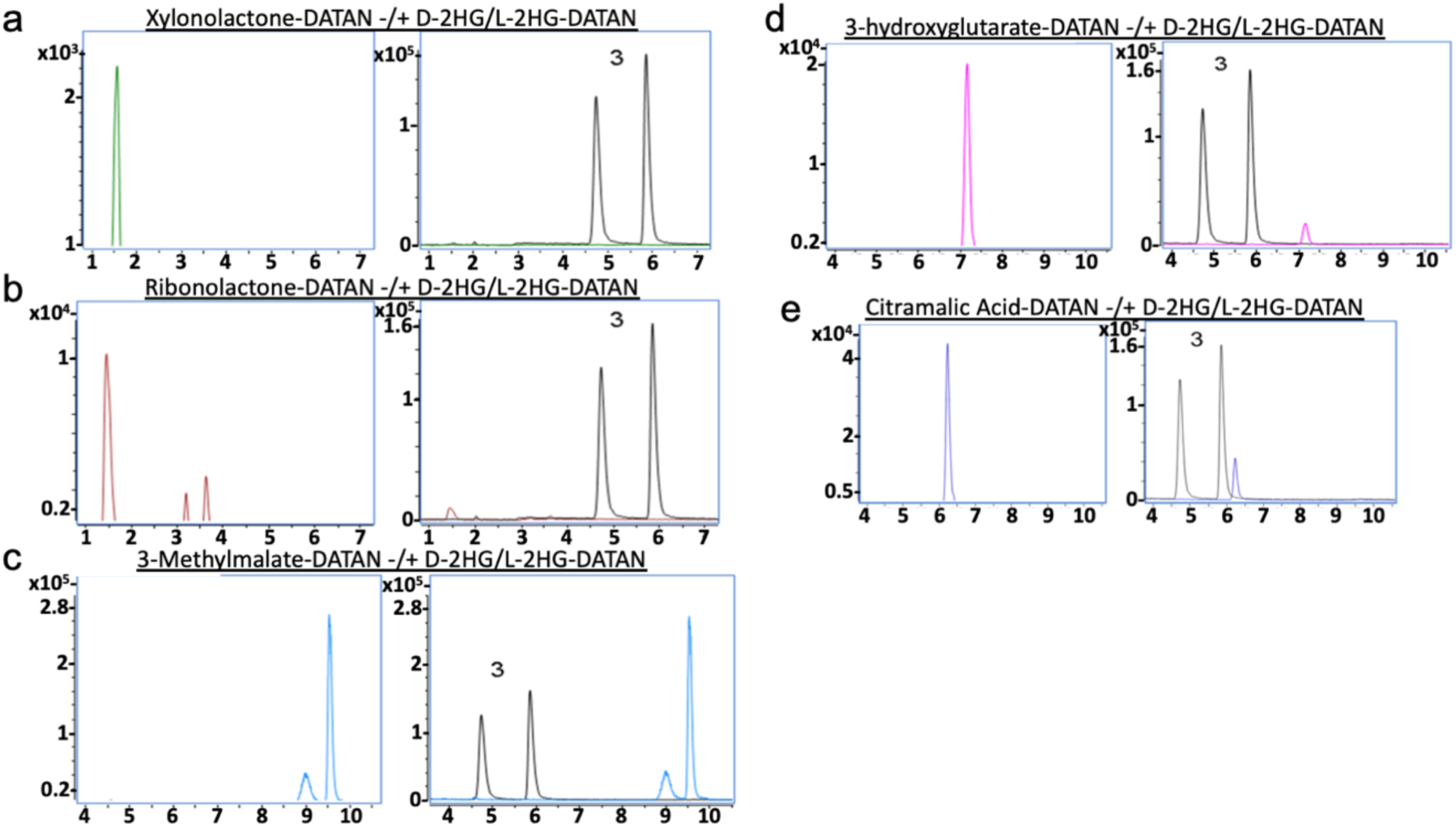
Extracted ion chromatograms (XICs) at m/z 147.0299 ± 5 ppm for L-2HG and D-2HG (3-black, left to right) along with the analytes defined as endogenous structural isomers with the identical mass for derivatized standards (3a-e). a) Xylonolactone (green); b) Ribonolactone (brown); c) 3-Methylmalate (light blue); d) 3-Hydroxyglutrate (magenta); e) Citramalic acid (dark blue). a) to e) Chiral derivatization with DATAN was applied to proper standards prepared at 26.0 nmol/mL for each 2-HG isomer, then each standard analyzed according to the proposed methodology for D-2HG and L-2HG quantification. Standards were prepared at 26 nmol/mL. Standards were injected at 6µL as indicated in Methods section. Abundance in counts per sec (y-axis) and acquisition time in minutes (x-axis) were omitted for clarity.

Commercial standard of known endogenous structural isomers of 2-hydroxyglutarate (m/z 147.030) were analyzed and compared to confirm that each isomer can be distinguished from D-2HG and L-2HG at separate retention time following derivatization with DATAN. In addition, the same standards were prepared without chiral derivatization and injected together with D-2HG and L-2HG and XICs were overlaid to confirm that they are isobaric (Supplementary Figure S1). This representation illustrates that our optimized methodology enhanced the overall separation achieved between D-/L-2HG and other endogenous structural isomers with non-chiral liquid chromatography on the Zorbax Eclipse C18 column (Figure 3 and Supplementary Figure S1). Furthermore, a survey of MS data were examined for biological sample matrices (for cellular as well as mouse glioma and contralateral brain tissue) commonly utilized in our IDH1-mutant glioma study following DATAN esterification (Supplementary Figure S2). The findings for citramalic acid (CM), which is the structural isomer that elutes the closest to D-2HG, did not reveal any apparent overlapping peak at interfering levels above LLOQ or basal levels for IDH^wt^ (Supplementary Figure S2). Lastly, mass spectrum search for m/z 152.047 for U^13^C-D-2HG/L-2HG did not return any known endogenous isomers with similar mass ±5 mDa to the internal standard using the Human Metabolome Database [34,35].

### Linearity

Calibration curves generated using simple (unweighted, normal) linear regression allowed each point to be weighted equally while confirming dilution linearity with a reproducible, reliable L-2HG (R^2^ = 0.995) and D-2HG standard curve (R^2^ = 0.997) (Figure 4). The curve was generated via repetitious, duplicate injections of the same diluted racemic mixture of 2HG standards representing a linear range using standards prepared with the concentrations of 104, 73, 52, 42, 31, 21, 5, 4, 3, 2, 0.8 nmol/mL. A lower limit of quantitation (LLOQ) was determined at 0.8 nmol/mL using standard preparation and TOF-MS experimental conditions as described in the Method section.

**Figure 4.**
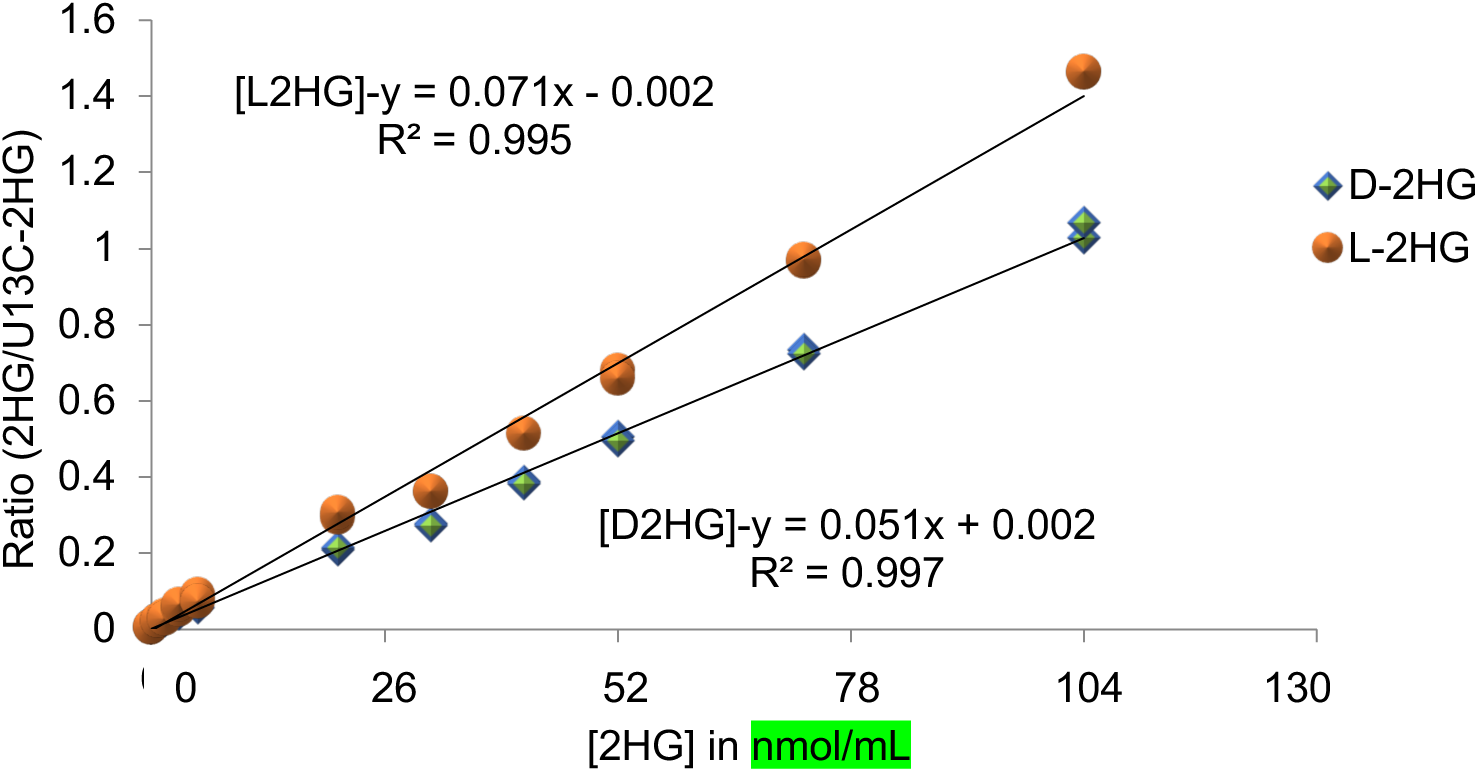
The representative calibration curves for D-2HG (blue) and L-2HG (red) base of the DATAN esters illustrate linearity over a range of 0.8-104 nmol/mL. The curves were produced from integrating the D-2HG and L-2HG area normalized to area of their respective internal standard, U^13^C-D-2HG and U^13^C-L-2HG. Each standards dilution was prepared in 70% methanol (aq) spiked with internal standard and dried under nitrogen gas stream. Dried standards were derivatized and analyzed via LC/MS with Zorbax Eclipse RRHD C18 column as described in Method section.

Previously published assays conducted on triple quadrupole (QQQ, or TQ) MS/MS instruments (e.g. Xevo TQ, API 3000 TQ Finnigan LTQ) with MRM capabilities exceed the sensitivity of single quadrupole instruments like the Agilent 6545 Q-TOF-MS system used for this assay; consequentially, comparatively higher LLOQ (i.e. less sensitivity on QTOF) is expected [28,29,31,36].

### D-2HG and L-2HG Peak Symmetry and Resolution

Given that both D-2HG and L-2HG elute under isocratic conditions at 3% solvent B, there are limitations to introduce further changes to the gradient in order to improve the peak chromatography. However, procedures were implemented to optimize the chromatographic conditions (such as, testing, comparing and adjusting solvent additives and composition) as detailed in the Method Comparison and Optimization section. As detailed below, the peak symmetry and resolution are reasonable.

The XICs shown in Figure 5 were used to determine the tailing factor (TF) for L-2HG (red) and D-2HG (green). Tailing factor is calculated at 5% from the baseline to the maximum peak height, as defined:

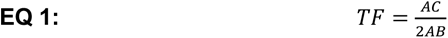

**Figure 5.**
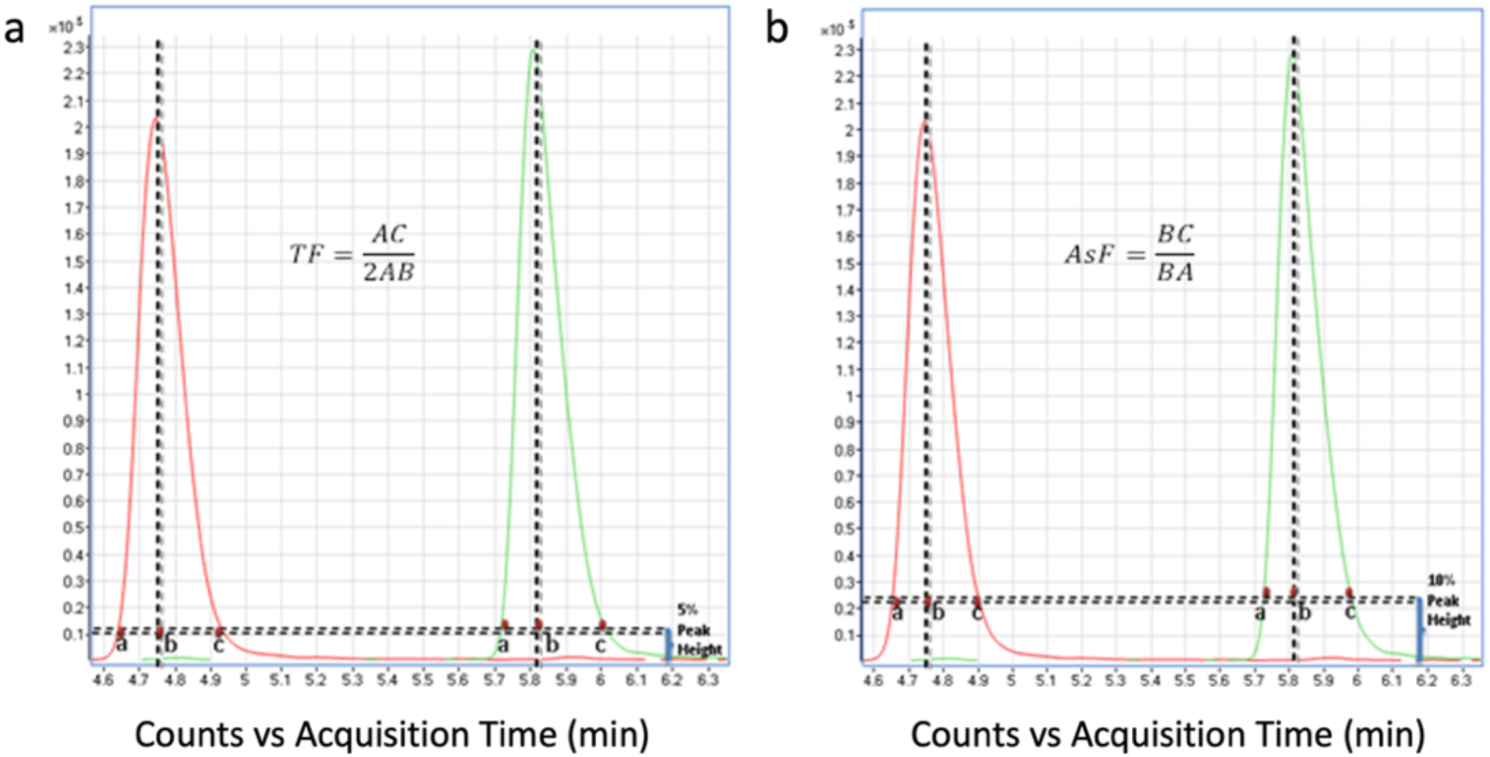
Schematic representation of tailing factor (TF) and asymmetry factor (AsF) assessment of D-2HG (green) and L-2HG (red) standards prepared at 26.0 nmol/mL, derivatized with DATAN, and analyzed via LC/MS as described in the Methods section. Intensity in counts per sec is represented (y-axis) and acquisition time (x-axis) are displayed for each XIC.

Whereas the peak width is measured in minutes from the upslope, *A* of the peak to the downslope, C and divided by twice the minutes between the peak maximum, *B* of the peak and point *A*. TF calculated for L-2HG (red) and D-2HG (green) was 0.520 and 0.516, respectively. A TF ≤ 2.0 is recommended for quantitative analysis involving high performance liquid chromatography (HPLC).

The asymmetry factor is calculated to quantify the peak tailing at 10% from baseline to the maximum peak height, as defined:

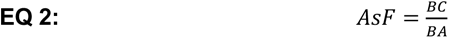

Similarly, the minutes from the peak maximum, B to the downslope, C is divided the minutes from the peak maximum, B to upslope A. AsF calculated for L-2HG (red) and D-2HG (green) was 1.05 and 1.10, respectively.

Peak resolution (*Rs*) was measured to quantify the separation of the peaks, as defined:

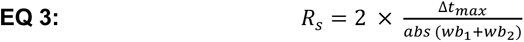

Where Δ*t_max_* indicates the time difference between the maximum peak heights (apexes) for the two peaks; *wb*_1_ indicates the baseline peak width for peak 1(L-2HG); and *wb*_2_ indicates the baseline peak width for peak 2 (D-2HG) [28,37]. Baseline separation between L-2HG and D-2HG presented an acceptable *R_s_* of 1.6 (at retention time of 4.75 min and 5.82 min, respectively).

### Precision and Accuracy

The same day and inter-day coefficient of variation (CV) % was measured to determine reproducibility using commercial standard quality control (QC) and pooled cell sample QC injections (Table 2).

**Table 2.** Intra- and inter-day precision for quantification of D-2HG and L-2HG standards by our method*.

| Analyte | Intra-day 1 | Intra-day 2 | Intra-day 3 | Inter-day | Calculated (sd) | Theoretical | Relative |
| --- | --- | --- | --- | --- | --- | --- | --- |
|  | CV% (n = 3 ) | CV% (n =5 ) | CV% (n =4 ) | CV% (n =12 ) | mg/L (n =12) | mg/L | error % |
| D-2HG | 5.3 | 5.0 | 1.5 | 4.0 | 3.94 (0.16) | 4 | -1.5 |
| L-2HG | 6.6 | 8.0 | 3.6 | 6.3 | 4.11 (0.26) | 4 | 2.7 |

average calculated concentration (n = 12) with standard deviation (sd) for both D-2HG and L-2HG was compared with the theoretical concentration in order to examine the accuracy of quantitation in terms relative error percent (%) across interval injections of commercial standard QC level over a 3-day cycle. Standards were prepared, derivatized and analyzed as described in Method section. All samples were maintained under isothermal conditions (4°C) in the autosampler over the course of the experiment.

Table 2 above provides the intra-day and inter-day precision findings for interval injections of the pure standard QC containing racemic mixture of D-2HG/L-2HG derivatized with DATAN that were then measured via LC/MS analysis. Additional results are provided for the pooled biological sample QC (composed of human-derived brain tumor cells ± IDH1 mutation) in the Supplementary Table S1. Overall, the CV% for intra-day and inter-day variability at the commercial standard and pooled biological (tumor cells) sample QC levels were below 10%.

### Recovery and Carryover

Recovery of 2-HG enantiomers was assessed using biological replicates of wild-type GSC923 glioma spheroids represented as groups of non-spiked controls (n=3) and spiked test (n=3) with 13 nmol/mL racemic-2HG standard mix following cell lysis. Spheroid cells were grown, collected, extracted, derivatized and injected using the proposed methodology as described in Methods section.

While Figure 6a only provides an XIC overlay of enantioselective detection of D-2HG and L-2HG for technical replicates (n = 2) of rac-2HG standard QC, the XIC overlays, Figure 6b allow us to compare the detection of basal, intracellular D-2HG and L-2HG levels in non-spiked (green) to D-2HG/L-2HG spiked (red/magenta) biological replicates of IDH1^wt^ GSC923 neurosphere extracts following chiral derivatization with DATAN.

**Figure 6.**
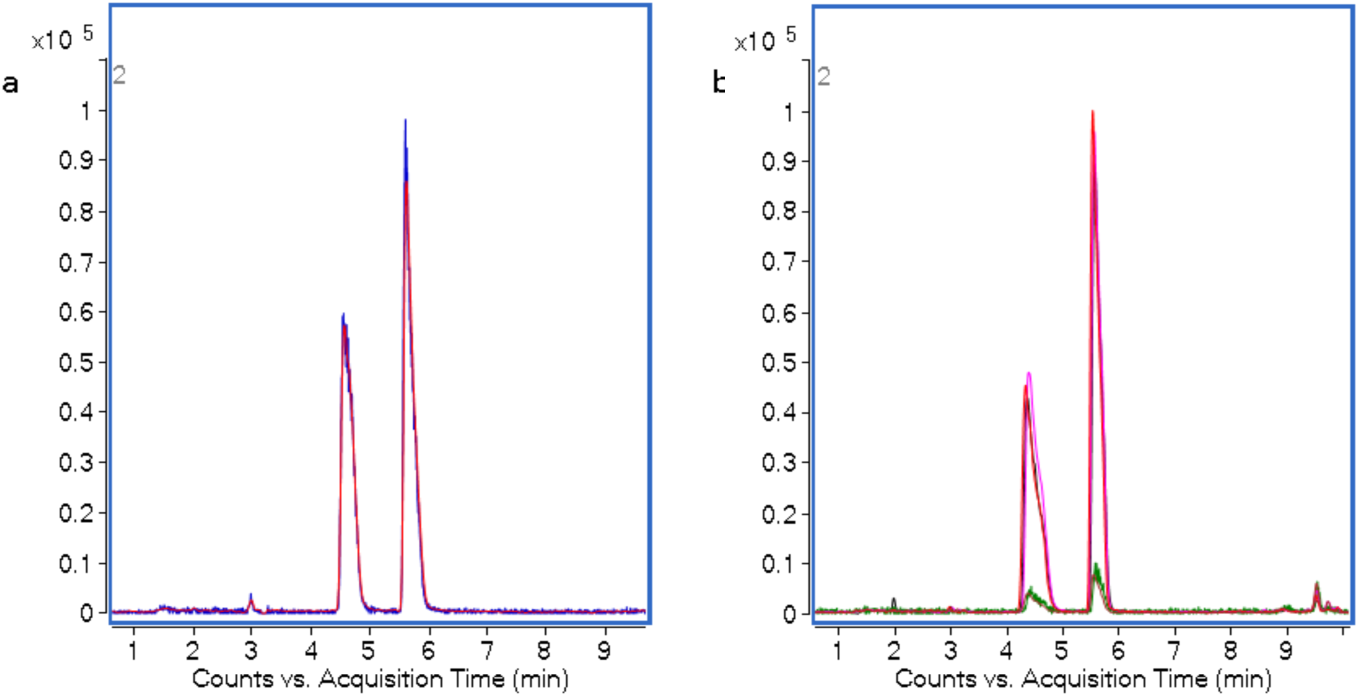
Extracted ion chromatograms (XIC) reveals the overlap between the biological replicates of wild-type IDH1, GSC923 glioma cell line. Chromatogram in a) shows the XIC overlay of duplicate injections of derivatized standard at 13 nmol/mL D-2HG and L-2HG (n=2) while b) shows the XIC overlay of non-spiked (green; n=3) and D-2HG/L-2HG spiked (red/magenta; n=3) replicates. Dried standards and samples were prepared, resuspended in 0.1 mL H2O and analyzed via LC/MS following conditions defined in the Methods section. Abundance in counts per sec is represented (y-axis) and acquisition time (x-axis) are displayed for each XIC.

Based on the data from Table 3, the target quantification of 2-HG enantiomers presented quantitative accuracy > 96% (n=3) and recovery > 90% (n=3) for the detection of D-2HG and L-2HG in sample matrices using the proposed MS-based analysis. The table also reveals that accuracy of quantitation exceeds 96% for D-2HG; wherein L-2HG quantitation was slightly above 100% accuracy indicating that calculated concentration L-2HG slightly exceeded the expected while remaining within reasonable range expected for relative error +2.7% (as defined in the Table 2).

**Table 3.** Recovery data for D-2HG and L-2HG quantitation with the proposed LC-MS method*.

| Spiked Analyte | Spiked Amount (nmol/mL) | Mean Concentration (nmol/mL) | CV (%) | Accuracy (%) | Recovery (%) |
| --- | --- | --- | --- | --- | --- |
| L-2HG (n=3) | 0 | 2.05 (0.11) | 5.22 |  |  |
|  | 13 | 15.28(0.29) | 1.91 | 101.8 | 90.6 |
| D-2HG (n=3) | 0 | 2.12(0.13) | 6.1 |  |  |
|  | 13 | 14.61(0.24) | 1.61 | 96.1 | 94.1 |

\*IDH-wt GSC923 neurospheres were individually spiked with 13 nmol/mL of each analyte (D-2HG and L-2HG). Following sample preparation and derivatization, dried samples and standards were resuspended in 0.1 mL H<sub>2</sub>O and analyzed as described in Method section. Accuracy was determined for mean concentrations (n=3) calculated after the area of each analyte peak (m/z 147.030) was normalized their respective internal standard, U<sup>13</sup>C-D-2HG and U<sup>13</sup>C-L-2HG (m/z 152.047). Recovery was determined for each analyte based on the mean area in pre-spiked sample – area of non-spiked sample extracts compared to the area detected for pure analyte standard prepared in 70% MeOH (aq) followed derivatization and analysis as described in Method section.

Moreover, negligible carryover (<1%) was observed in the blank following injection of high concentration of 73 nmol/mL racemic 2HG (rac-2HG) standard. This observation was based on the peak area detected in for D-2HG/L-2HG-analog (m/z 147.030) and a D-/L-2HG-DATAN esters (m/z 363.057) at the elution times corresponding for each enantiomer following derivatization with DATAN and TOF-MS analysis (Supplementary Figure S3).

### Degradation

Degradation of 2HG-DATAN esters was assessed based on the percentage of underivatized 2HG analog area detected compared to the area sum of total racemic 2HG detected in daily duplicate injections of the rac-2HG standard that was prepared, derivatized and analyzed as described in the Methods section.

The data in Table 4 revealed that degradation can occur at a rate ≤ 0.05%/day for 2HG-DATAN esters and respective internal standards in racemic (rac) standard mixture containing 21 nmol/mL D-2HG and L-2HG (m/z 147.030) and internal standards at 9.8 nmol/mL U^13^C-D-2HG and U^13^C-L-2HG (m/z 152.047) reconstituted in 100 µL of H_2_O (LC/MS grade) over 3-day period at 4°C. In addition to the low degradation percentage (%) with standard deviation (sd) shown (Table 3), the low relative error % reported (Table 2) for periodic injections of rac-2HG standard QCs revealed that the percentage of deviation between the measured concentrations was consistently less than 10% from the expected concentration during the 3-day assessment; a metric used to assess stability in a previous publication [29].

**Table 4.** Time-dependent degradation of 2HG-DATAN esters over 3-days.

| Analyte | Total 2-HG Degradation (sd) |  |  |
| --- | --- | --- | --- |
|  | Day 1<br>avg % (n = 2) | Day 2<br>avg % (n = 2) | Day 3<br>avg % (n = 2) |
| DL-2HG | 0.008 (0.011) | 0.098 (0.005) | 0.150 (0.001) |
| U13C-DL-2HG (IS) | 0.016 (.003) | 0.093 (0.008) | 0.144 (0.016) |

### Matrix effect

A matrix factor of +1.2 was determined for L-2HG and -0.9 was determined for D-2HG for spiked in the pooled sample QC composed of wild-types (GSC923) and IDH1 mutant (TS603) samples (Supplementary Table S6).

### Method Comparison and Optimization

Assessment of published applications of enantiomeric quantification of 2HG [28,30,31,36] and in-house alterations were conducted to define reproducible derivatization and to optimize LC conditions for the detection and separation of both 2-HG enantiomers.

### Comparison of Derivatization Method

In order to select a robust approach for chiral derivatization of 2-HG enantiomers, we explored a set of applications and compared stability and reaction completion following esterification with D-2HG/L-2HG with DATAN and TSPC. The first step was to determine whether chiral derivatization with DATAN was a suitable approach to facilitate differentiation of 2HG enantiomers. For this step, multiple published and in-house approaches, detailed in Supplementary Table S2 were introduced to optimize our approach for high-throughput applications [28,30,31,36]. Based on our assessment, consistent and stable derivatization was achieved by adding 60 µL of 230 mM DATAN in 1:4 acetic acid/dichloromethane (DCM) to each sample (without catalysts) followed by mixing (1400 rpm) at 75 °C for 32 min; in addition to drying derivatized samples (∼2h) at 35°C with speed vac concentrator (Figure 6).

While chiral derivatization with DATAN facilitates direct detection and quantification of the 2-HG moiety (m/z 147.030) from the DATAN ester (Figure 2), derivatization with TSPC relies on the quantification of the intact ester, 2HG-TSP (m/z 448.108) in negative ion electrospray mode (ESI-) as shown in Figure 7a (below). The main benefit of the TSPC esterification remains the potential to reach chemical equilibrium with 2HG using a nominal concentration ranging from 0.74 to 1.25 mM TSPC derivatizing agent in acetonitrile. In addition, TSPC reagent (100 µL) and pyridine (2 µL) was added under mild conditions (incubation for 10 min and drying via N_2_ gas concentrator at 25 °C) which can preserve labile analytes.

**Figure 7.**
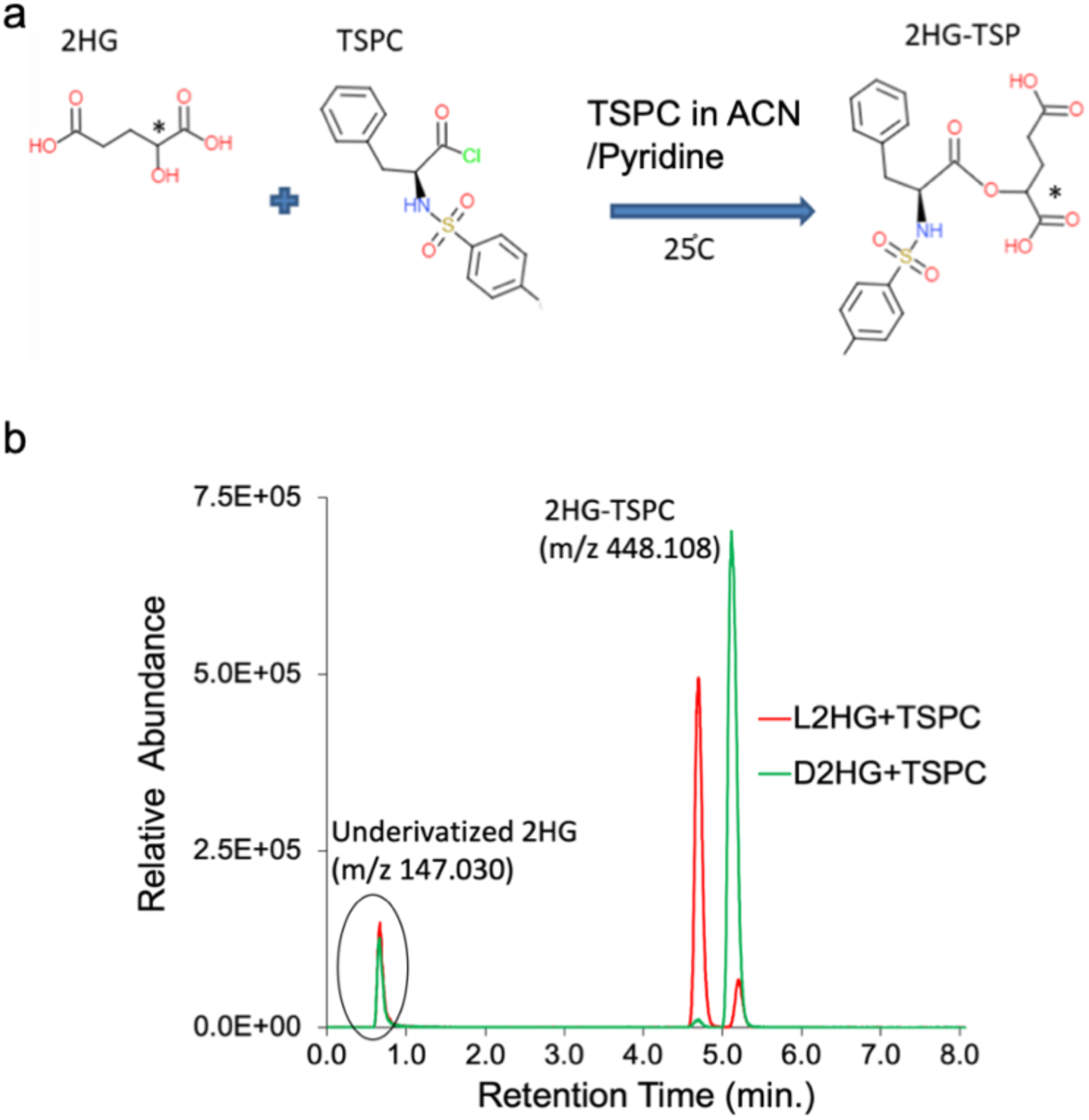
Chiral derivatization of 2-HG racemic with TSPC. a) The esterification reaction of 2HG with TSPC to form 2HG-TSP. b) XIC reveals high resolution in the detection of 52 nmol/mL D2-HG and L-2HG following derivatization with TSPC and subsequent separation on BEH Phenyl C18 2.1x50mm column over 9 min gradient with mobile phases A (5mM ammonium acetate in water) and B (100% methanol). Highlighted in the green circle is the amount of racemic 2HG that elutes below 1 min and represents approximatively 19% of the total compounds. L2HG and D-2HG are represented in red and black respectively. Relative abundance was determined in counts per sec (y-axis).

The chromatogram in Figure 7b demonstrates that optimal separation was acquired on the Xbridge BEH Phenyl C18 (Waters Corporation, Milford, MA, USA) following chiral derivatization of 2HG with TSPC at 25°C for 10 min. The assay achieved an ideal resolution, R_s_ = 2.1 between D-2HG and L-2HG when applying EQ 3 (as described in Section 2.2.3).

A true comparison between the two similar approaches was done in prior methods which relied on the detection and quantification of 2HG-DATAN esters (m/z 363.057) account for ionization efficiency and 100-fold lower sensitivity than for 2HG-TSP esters (m/z 448.108) [28,29,30]. However, in contrast to the methods, our analysis relies exclusively on the enantioselective detection of D-2HG and L-2HG analogs (m/z 147.030) using TOF-MS with collision energy of 0V. Therefore, this type of comparison was not conducted here.

In comparison to the DATAN esterification, the assay that relied on TSPC esterification exhibited higher resolution with relatively similar retention and detection of 2HG enantiomers. However, unlike DATAN, TSPC requires special storage conditions under pressurized argon for preservation. It was inferred that TSPC can be more reactive; thus, less stable under standard storage and experimental conditions. At the onset of analysis, a substantial amount of degradation product present in the form underivatized 2HG standard (≈19% D-2HG/L-2HG) was detected via LC/MS following esterification with TSPC reagent (Figure 7b). These findings were attributed to the high hygroscopicity and related degradation of the TSPC. In contrast, 2HG-DATAN esterification consistently presented derivatization that exceeded 99% of the total detectable 2-HG (Figure 6) using our proposed methodology as detailed in Methods section. Furthermore, 2HG-DATAN esters presented stability exceeding 72h; at which point underivatized 2HG presented in the sample was found to be 0.15% of the total 2HG (Table 4). Further optimization of TSPC-based high-throughput 2HG quantification was abandoned due to challenges to maintain stable 2HG-TSP esters and incomplete derivatization of analytes. In our workflows, stability is paramount for high-throughput analysis involving large scale experiments that extend for multiple days.

### Liquid Chromatography Optimization

The second step was to evaluate different LC applications in order to identify an analytical column and combination of reagents that demonstrates well-retained, well-resolved, and reliable MS detection of 2HG enantiomers. A set of liquid chromatography conditions (provided in Supplementary Table S3) were explored to optimize the separation, retention and detection of the 2HG enantiomers. Although, it should be noted that particle size and length of column are contributing factors, these variables were not compared. Columns having C18 and phenyl stationary phases were initially assessed using the reverse phase gradient conditions described in Supplementary Table S3. The initial assessment was based on the use of Xselect HSS T3 (High Strength Silica with proprietary endcapping) column to separate and detect 2HG enantiomers following derivatization with DATAN, as previously published [38].

Zorbax Eclipse Plus RRHD (Rapid Resolution High Definition) provided enhanced retention, detection, separation and peak quality under the same gradient condition compared with the other columns tested. After determining that substitution of Xselect HSS T3 with Zorbax Eclipse Plus RRHD C18 column improved the chromatography for 2HG enantiomers (Supplementary Figure S4). Upon comparison of HILIC columns using HILIC method described for Supplementary Figures S5 and S6, we did not observe an added benefit to incorporating HILIC conditions and chiral derivatization with DATAN (as described in Method section) to enhance enantiomeric selection of 2HG analogs. Additional steps were taken to optimize other LC conditions (including adjustments in composition of gradient buffers and additive, diluent) detailed in Supplementary Table S3. Herein, we present the combination of mobile phase buffers, diluent and buffer additives that greatly improved the chromatographic separation and detection of 2HG enantiomers.

Based on our assessment of optimized LC conditions, an in-house alteration in composition of organic mobile phase to 95:5 MeOH/ACN (in place of MeOH only as previously published [6]), reduced peak broadening and improved analyte retention (Supplementary Figure S7a). For the optimized LC conditions, aqueous mobile phase was also composed of H_2_O containing 2 mM ammonium acetate (NH_4_Ac) buffer and formic acid titrated to pH 3.5 with caution. After determining the optimal column and liquid gradient compositions, an assessment of various diluents (listed in Supplementary Table S3) initially revealed that reconstitution in 50% ACN in aqueous (aq) solution provided good retention. Follow-on comparison with reconstitution in 100% water revealed improved detection without an apparent compromise in stability of 2HG-esters; likewise, there was not an apparent increase of degradation products (underivatized 2HG) in the XIC (Supplementary Figure S7b). Our assessment revealed that chromatography and detection of 2HG enantiomers was notably enhanced when buffer additive, ammonium formate (NH_4_HCOO) was substituted with ammonium acetate (NH_4_Ac) buffer (Supplementary Figure S7c); an adaption to previously published assays [39,40].

### Detection of Enantiomeric Impurities in D-2HG and L-2HG Standards

During the method optimization, there were challenges to identify a source for high purity 2HG standards. In order to resolve these concerns, comparisons were conducted using consecutive injections of D-2HG and L-2HG standards from two different vendors under the same experimental conditions. In the development of the assay, it was found that multiple L-2HG standards acquired from an initial distributor were all suspected of being contaminated with D-2HG. The purity issue would have been undetected without the use of chiral derivatization.

Figure 8 provides XICs from the 2-HG standard after injecting (26 nmol/mL) to examine chromatographic symmetry (in Section 2.2.3), detection and purity of D-2HG/L-2HG standard. The difference in peak intensity using racemic-2HG standard suggests that L-2HG consistently presents lower peak intensity than D-2HG at the same concentration under the assay conditions. Despite differences in ionization, accurate quantification of L-2HG in biological samples is expected using proposed MS-based detection method based; this is evident given that both, L-2HG and D-2HG standards were assessed with a recovery >90% (Table 3). Figure 8b reveals that the challenge of determining which commercial standards present enantiomeric impurities can be resolved via DATAN in order to improve reliability of enantioselective 2HG quantification. Analysis of L-2HG (red) and D-2HG (green) moiety of the 2HG-DATAN esters confirmed that the initial L-2HG commercial standard contained an impurity (∼12% of the total 2HG) detected at m/z 147.030. The retention time and mass for the impurity was consistent with D-2HG. This finding suggests that the L-2HG purity was below the purity ≥ 98% reported by the distributor label. Whereas XICs from the analytical grade D-2HG and L-2HG standards from an alternate distributor, Millipore-Sigma (Figure 8c) clearly surpassed the manufacturer’s reported purity ≥ 98%. As a result, the proposed methodology was optimized to only use the analytical grade D-2HG and L-2HG. This outcome illustrates how the application of chiral derivatization provides quality assurance via LC-MS enhanced detection and measurement of standards contaminated with enantiomers.

**Figure 8.**
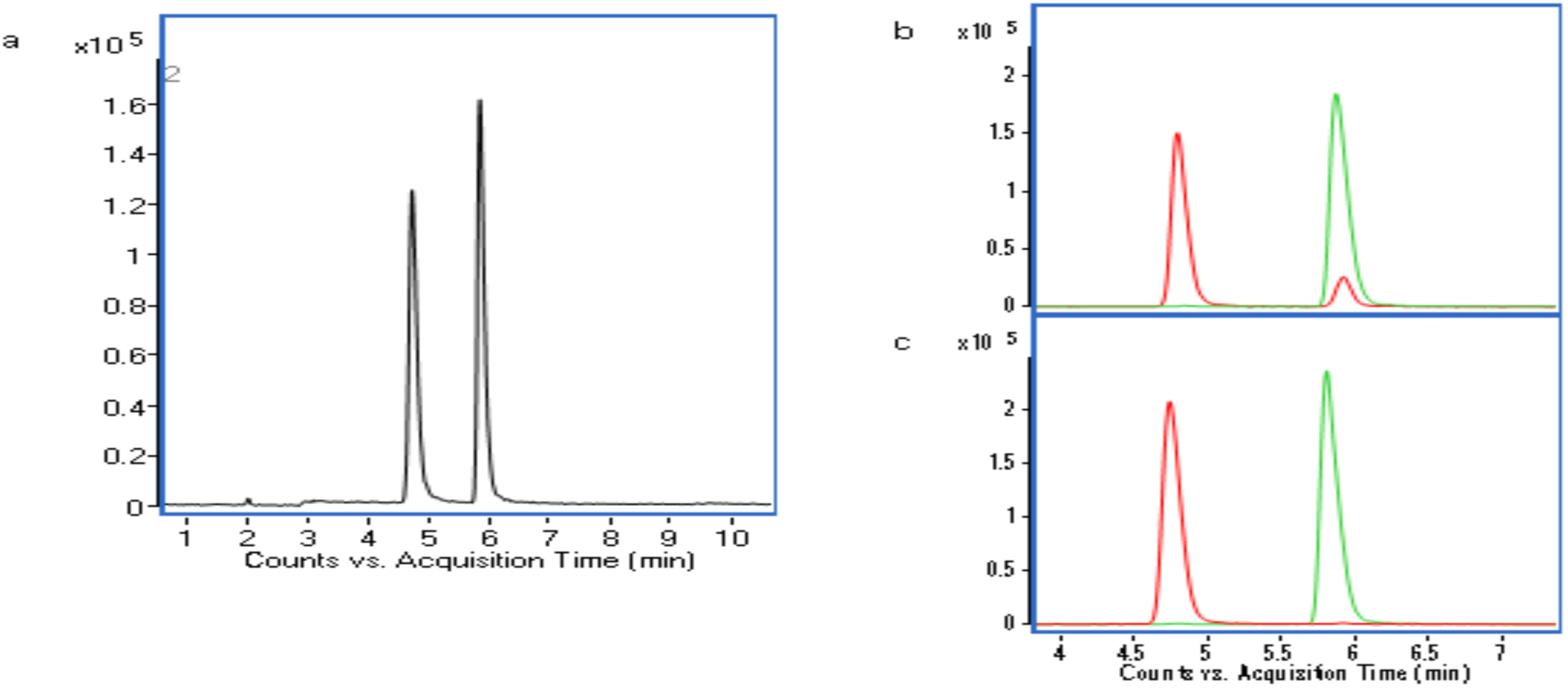
The XICs reveal a) differences in peak intensity for L-2HG and D-2HG using DL-2HG standard which contains a racemic mixture of D-2HG and L-2HG; b) XIC for commercial standards of L-2HG (red) and D-2HG (green) purchased from initial supplier for which L-2HG standard present enantiomeric impurity that corresponds to D-2HG; c) analytical grade L-2HG (red) and D-2HG (green) from Millipore Sigma demonstrate ideal purity. The abundance is represented as counts per sec (y-axis) and the acquisition time is represented in minutes (x-axis). Standards were prepared at a concentration of 26 nmol/mL, derivatized with DATAN and analyzed via LC/MS as detailed in method section.

### D-2HG and L-2HG quantification in biological specimens

Following our optimization of the proposed LC/MS assay for D-2HG/L-2HG quantification, lastly, we applied to methodology to disease-relevant biological samples.

Figure 9a-c illustrates the D-2HG/L-2HG concentrations in various human-derived glioma samples including: heterozygous (IDH1^mut^/^wt^) spheroid TS603 (Oligodendroglioma III), and NCH612 (Anaplastic Oligodendroglioma III) glioblastoma multiforme (GBM, IDH1^wt^) including adherent cells U251 and spheroid GSC923; xenograph mice with intracranial mutant IDH1 glioma. Each specimen was analyzed following sample preparation and assay conditions described in the Methods section. Figure 9b suggests that D-2HG concentrations are substitution-dependent when IDH1^wt^ U251 cells with genetically engineered IDH1^R132H^ (Rh) are compared with IDH1^R132C^ (Rc). U251^R123H/C^ genetically-engineered to express IDH1 mutation entail a substitution of arginine at codon 132 (Arg132) with histidine (R132H) or cysteine (R132C) in IDH1 gene (which normally translates to a docking site for isocitrate in wild-type IDH1); alternatively, increasing affinity for NADPH and catalyzing neomorphic reduction of 2-OG to D-2HG in mutant IDH1 [8] [42]. Figure 9b suggests that the R132H allele (occurring in ∼90% of the IDH mutant gliomas) can be more efficient in converting 2-OG to D-2HG than R132C [43]. Figure 9c demonstrates that a peripheral amount of D-2HG can be detected in nontumor regions for xenograph mice with intracranial IDH1 mutant glioma. This suggests that D-2HG is exported from mutant IDH1 gliomas and infiltrates the nontumor regions brain which has implications for the tumor microenvironment. 2HG stereoisomers were detected in both cells (Figure 9d) and media (Figure 9e). Interestingly, more D-2HG is exported to the microenvironment than retained in the representative IDH1 mutant glioma, TS603 cell line. Figure 9f demonstrates that the total cellular D-2HG/L-2HG production (based on combined cell and media concentrations) was not greatly impacted following 72h exposure to hypoxia (3% O_2_) compared to normoxia. Figure 9g confirms that the total production and export of D-2HG (detected in media and cells combined) was substantially, albeit not completely, suppressed following treatment with 25 µM AGI-5198.

**Figure 9.**
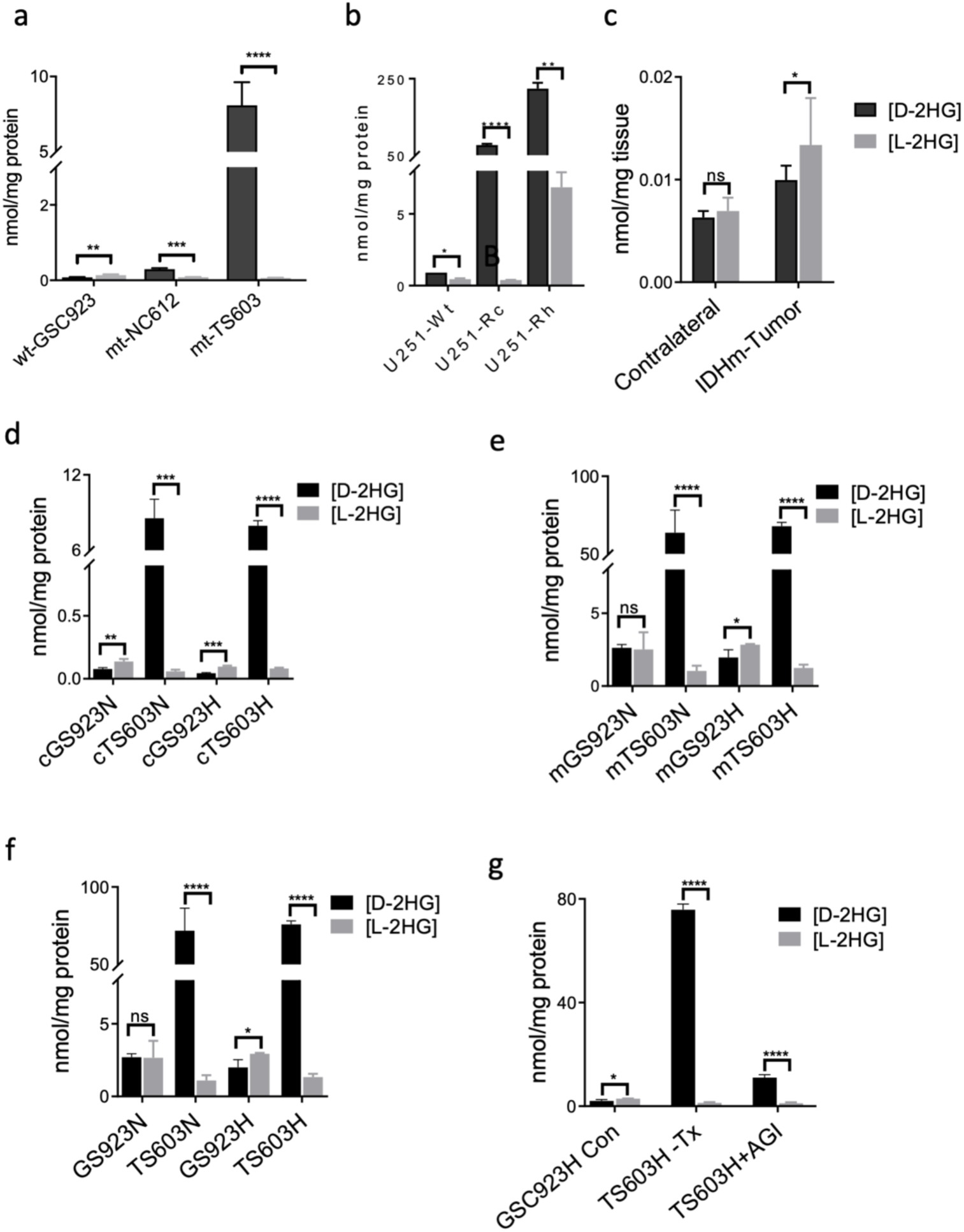
Quantification of D-2HG and L-2HG in nmol/mg protein for cells and tissues based on proposed LC/MS quantification methodology. a) D-2HG/L-2HG in stem-like spheroid cell lines: IDH1^wt/wt^ GSC923 (n=3), heterozygous IDH1^mut/wt^ NC612 (n=3) and TS603 (n=4) b) D-2HG/L-2HG in IDH1 ^wt^ and genetically engineered IDH1^R132H/C^ U251 cells (n=3). c) D-2HG/L-2HG in mutant IDH1 ^R132H^ gliomas and contralateral (nontumor) brain extracted from xenograph mice (n=3)[41]. Comparison of D-2HG/L-2HG concentration under normoxic (N) and hypoxic (H) conditions d) for cells only; e) exported to corresponding media only; f) for media and cells combined under H and N for representative IDH1^wt^ GSC923 and IDH1^mut^, TS603; g) following treatment (Tx) with IDH1 inhibition, AGI5198 (+AGI) IDH1 in IDH1^mut^ compared to baseline 2HG in control (Con, GBM). All samples were prepared, derivatized and analyzed via LC/MS as detailed in Method section. Statistical analyses were performed in GraphPad Prism 7.05 with Bonferroni-Dunn multiple test comparison for D-2HG and L-2HG concentrations within each sample. Error bars represent standard deviation between biological replicates. P-value are indicated as: ns, not significant; *, p≤ 0.05; **p≤ 0.005; ***p≤ 0.0005; ****p≤ 0.0001.

### Metabolic dysregulation that correlates with D-2HG accumulation

In addition to resolving and independently quantifying D-2HG/L-2HG, the proposed LC/MS methodology was used to conduct estimated relative quantitation and comparative analysis on central carbon metabolites (whose identities were validated by MS/MS and retention of standard to prepared a targeted extraction library) that are known to regulate oncogenesis and neoplasticity [44–47].

Figure 10 reveals the coverage of central carbon metabolites whose identities were validated based on retention times and tandem MS (MS/MS) fragmentation with pure standards (Supplementary Table S4). Relative quantitation and comparative analyses were implemented to examine the value and biological relevance of incorporating acquisition of key metabolites in conjunction with resolving and quantifying 2HG enantiomers.

**Figure 10.**
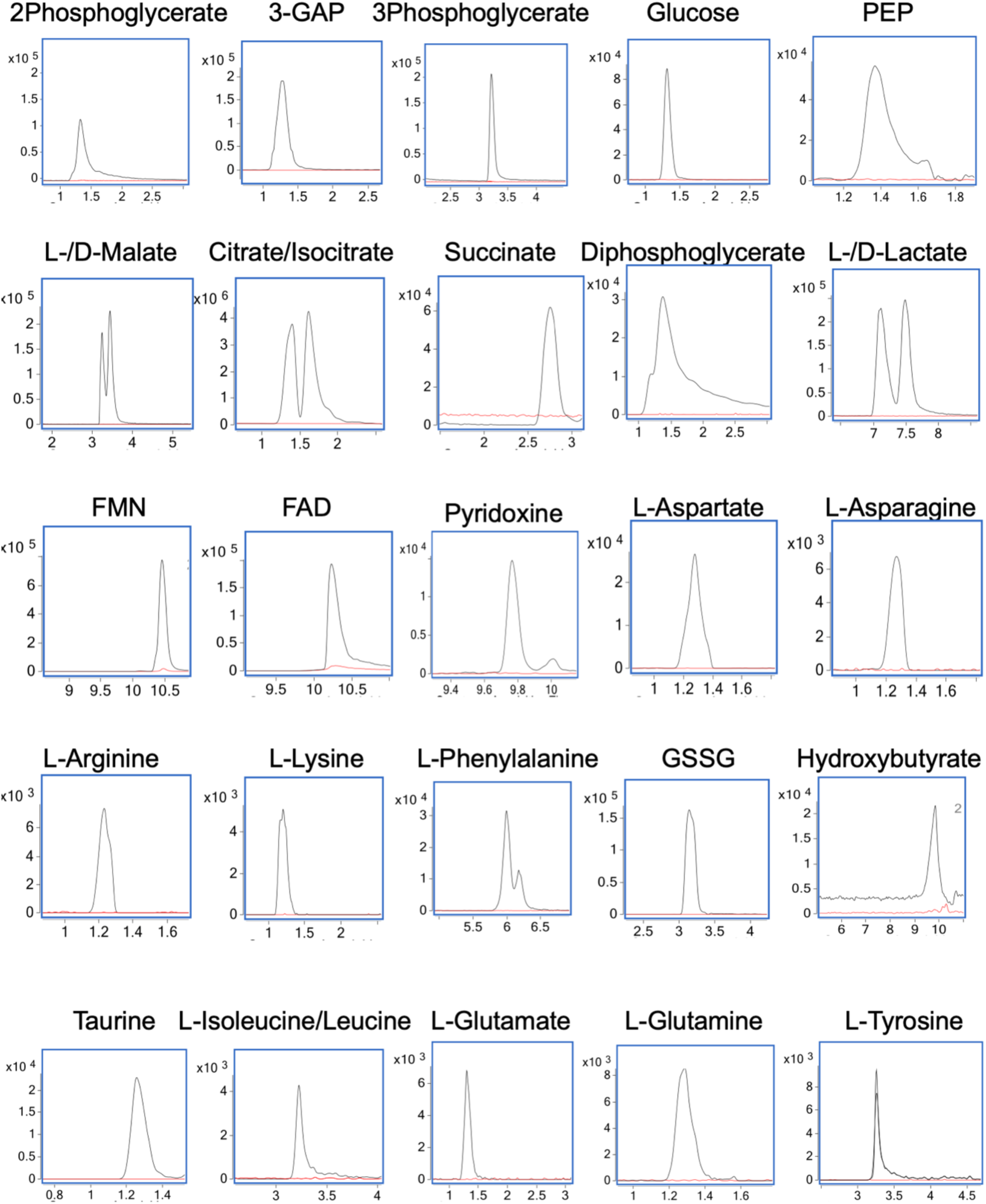
TOF-MS Detection and confirmation of key metabolites with standards. Commercial standards were prepared at 52 nmol/mL, derivatized with DATAN and analyzed by LC/MS conditions outlined in the Method section. The XIC for each analyte ((black) is shown overlaying their respective blank (red). The y-axis represents the relative abundance measured in counts per second (cps) and the x-axis represents retention time in minutes for which axis labels were omitted for space considerations. MS/MS–based validation was applied to produce transitions shown in the Supplementary Table S4, which were generated using the LC/MS/MS conditions outlined in the Method section and collision-induced dissociation (CID) by ramping the collision energy (CE) from 0 – 10 V. Commercial standards were purchased from Sigma Aldrich. Legend: 3-GAP, Glyceraldehyde 3-phoshosphate; FMN, Flavin mononucleotide; G1,3-P, 1,3-Bisphosphoglycerate; GSSG, Glutathione disulfide; PEP, Phosphoenolpyruvate. Only the L form of Phe, Glu, Gln, Arg, Asp, Asn, Lys, Tyr, Ile, Leu was investigated.

Comparative analyses were conducted for cells prepared under hypoxic (H) and normoxic (N) condition using the proposed methodology. Heterozygous IDH1*^mut^* TS603 glioma spheroids presented the highest innate production of D-2HG; thus, TS603 served as a representative to evaluate downstream metabolic changes attributed to elevated D-2HG concentration in mutant IDH1 gliomas. Homozygous IDH^wt^ GSC923 glioma spheroids were selected to represent wild-type metabolism given that GSC923 present the lowest cellular concentrations of D-2HG and a spheroid morphology that resembles IDH1^mut^ gliomas, such as TS603.

The main differences were observed for the comparison between representatives IDH1*^wt^* (GSC923, n = 3) and IDH1*^mut^* (TS603, n = 3) cell lines (Figure 11). As expected, the most significant change is the 106-fold elevation of D-2HG in the representative IDH1*^mut^*, TS603 when compared to IDH1*^wt^* GSC923 cells. A significant decrease in glutamine was evident in representative IDH1*^mut^* compared to IDH1*^wt^* cells (Figure 11), which is consistent with glutamine-dependency and consequent depletion to compensate oxoglutarate demands in IDH1*^mut^* gliomas [48,49]. Representative IDH1*^wt^* presented significantly elevated L-2HG compared to representative IDH1*^mut^* (Supplementary Table, S5), as anticipated. In relation, lactate was significantly elevated while succinate was significantly downregulated in IDH1*^wt^* compared to IDH1*^mut^* (Figure 11). Based on the elevated lactate (glycolytic end-product) and concomitant decrease in succinate (TCA cycle biomarker), it was inferred that Warburg effect is likely more prominent in the IDH1*^wt^* [50]. An imbalance between TCA intermediates and lactate production can occur when lactate dehydrogenase (LDH) activity prioritizes the conversion of pyruvate to lactate before it is shunted into TCA by PDC; moreover, HIF1α-driven overexpression of pyruvate dehydrogenase kinase (PDK) can deactivate PDC, thus hindering shunting of pyruvate to supplement TCA [51,52]. Elevated lactate was also found to inhibit oxoglutarate-dependent prolylhydroxylase, EGLN1 and activate HIF-1α in normoxic tumors [53].

**Figure 11.**
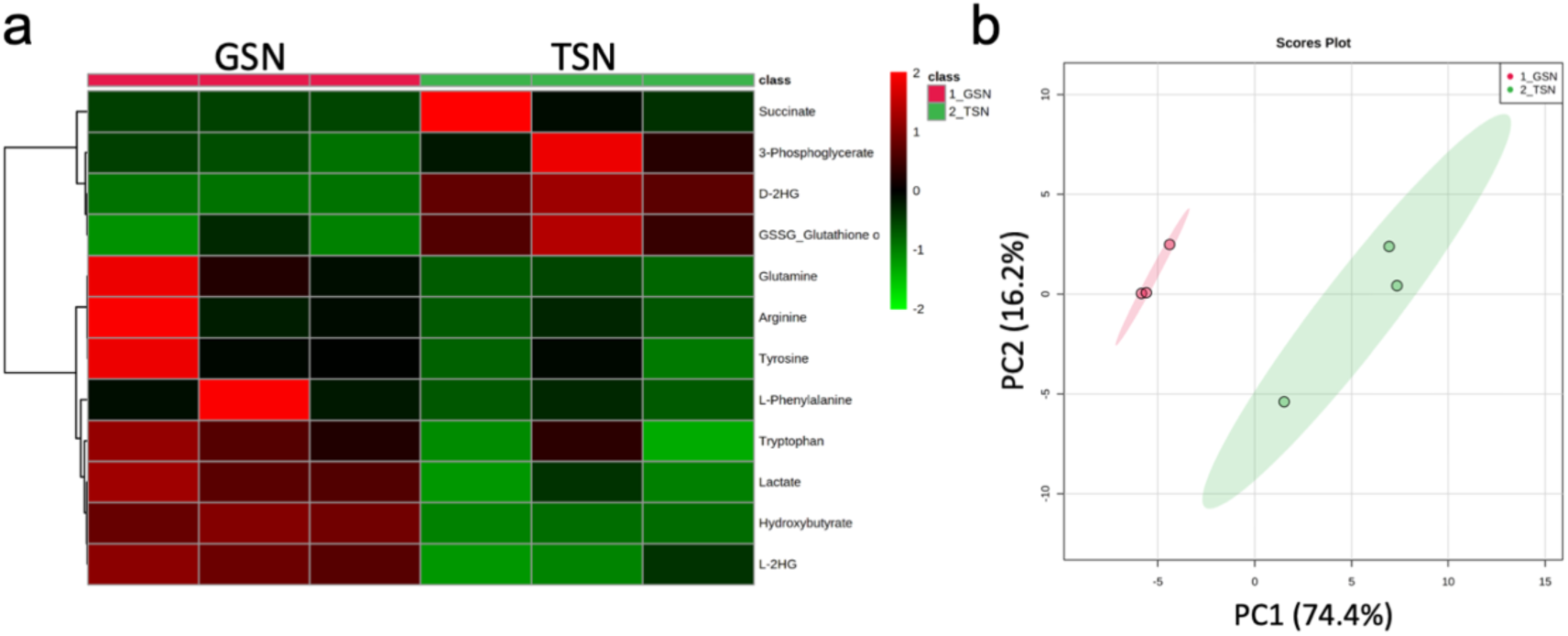
Metabolic analysis with key biomarkers of central carbon and cancer metabolism for IDH1*^mut^* TS603 (TSN) cells compared to IDH1*^wt^* GSC923 (GSN) grown under normoxic conditions, prepared and analyzed as described in Methods section. a). Heat map containing twelve most significantly altered metabolites between biological replicates of IDH^wt^ (red) and IDH^mut^ (TS603) (n=3). b). PCA plot shows discrimination between the two groups and was generated using MetaboAnalyst [35].

## Discussion

Reliability and reproducibility take precedence for bioanalytical applications utilized in high-throughput and clinical studies. One aim of this study was to examine and resolve challenges that can occur during the preparation of derivatized samples and enantioselective LC/MS quantification of 2-HG. The next goal was to propose a comprehensive methodology that defines a unique combination of sample preparation, derivatization and LC/MS applications that were selected and optimized to reduce foreseeable challenges in high-throughput, enantioselective quantification of 2-HG. Herein, we reported a reliable workflow that facilitates consistent derivatization, stability and separation of the 2HG enantiomers. Ultimately, implementation of the optimized methodology allowed characterization of D-2HG and L-2HG levels based on IDH1 mutation, cell-type, sample type and oxygen levels.

While several reagents (e.g., combination of DATAN, DCM and acetic acid) were conserved from previously published methods, our enantioselective methodology proposed an adapted application of the Zorbax Eclipse RRHD column and incorporated a unique combination of mobile phase composition and sample preparation techniques to conduct accurate quantification of D-2HG/L-2HG using MS-based detection. MS-based detection allowed for the added advantage of multi-target quantitation on the Agilent 6545 QTOF LC/MS instrument. Based on our assessment of the proposed LC/MS quantitative analysis for D-/L-enantiomers of 2-HG, the methodology demonstrated reasonable combination of stability, sensitivity and resolution to detect and differentiate between the 2-HG stereoisomers. In addition, the proposed assay facilitated the detection and differentiation of endogenous 2-HG isomers and biomarkers in central carbon metabolism.

Independent quantification of D-2HG and L-2HG achieved the following objectives: determined which cell lines derived from patients are the best representative models of the IDH1*^mut^* gliomas; examined the downstream advantages of metabolic reprogramming that upregulate the biosynthesis of either metabolite; and identified enantiomeric impurities in commercially available standards.

Firstly, our optimized methodology was able to validate the use of TS603 cell line as a good model system of oligodendroglioma with IDH1 mutation via presence of large amounts of D-2HG. TS603 cell line presented substantially higher concentrations of D-2HG than NCH612 IDH1*^mut^* cells. Since loss of mutant allele has been reported and is encountered in progressed gliomas or as an adaptation to cell culture conditions, the amount of D-2HG produced by model systems that are recapitulating IDH1*^mut^* glioma is a good indicator that mutant allele is preserved with serial passaging [54,55]. In our hands, TS603 oligodendroglioma cell line displayed the highest D-2HG concentration out of all the patient-derived cell lines investigated, validating the use of TS603 cell line for future studies. In addition, we found that a 72h exposure to 3% oxygen had a mild impact on the production of L-2HG within the representative IDH1*^mut^* cells.

Secondly, the proposed LC/MS methodology facilitated simultaneous quantitation of other metabolites and elucidated a significant dysregulation in metabolism between IDH1*^wt^* and IDH*^mut^*. Based on our data, IDH*^mut^* presented concurrent downregulation of lactate and glutamine that is consistent with prior research [48,56]. We found that glutaminolysis occurs as an anaplerotic response along with consequent glutamine-driven oxidative phosphorylation, potentially to stimulate alternative source of energy production in response to the shunting of pyruvate away from TCA cycle in IDH1*^mut^* gliomas [48,56–58]. The incorporation of targeted analysis with central carbon and disease-relevant metabolites in conjunction with the D-2HG and L-2HG quantification allows metabolic reprogramming associated to elevation in D-2HG or L-2HG concentrations to be examined simultaneously.

Thirdly, our method was capable of identifying impurities in commercial standards. This finding has important implications. We often rely on vendors to provide us with pure standards when in fact the pure enantiomers can be contaminated within each other as shown herein. Considering the divergent roles of D-2HG and L-2HG, the presence of L-2HG enantiomer in a pure D-2HG standard could lead to mixed results in the altered pathways downstream when used in biological assays and limit our ability to identify potential mechanistic insights.

In our glioma studies, we are focused on understanding the relationship between metabolic differences between gliomas to the concentration of D-2HG, particularly, produced in mutant IDH1 model systems. A thorough characterization of differing D-2HG concentrations across the heterogeneous mutant IDH1 cell lines and mice models is fundamental to validate available model systems and then, hypotheses that relate to downstream metabolic changes and other epigenetic occurrences that correlate with D-2HG levels only. The purpose of such investigation is to ultimately elucidate points of vulnerability that are specific to mutant IDH gliomas. Given that L-2HG can be produced in response to acidic and hypoxic condition, it is important to separately measure the contribution of D-2HG versus the total 2-HG. This facilitates accurate characterization of the role that D-2HG serves in mutant IDH1 cancers. Standard nonchiral LC/MS methods do not effectively resolve the enantiomers of 2-HG. Therefore, our previous LC-MS metabolomic studies relied on the quantification and relative comparison of total 2-HG levels. In our effort to increase confidence in our characterization of the IDH1 mutant cells, D-2HG-specific epigenetic changes, and metabolic reprogramming, we introduced the use chiral-derivatization agents into our workflow.

## Materials and Methods

### Standard Materials

D-2-hydroxyglutaric acid and L-2-hydroxyglutaric acid (2HG) disodium salt, ^13^C_5_-D- and ^13^C_5_-L-2HG disodium salt, Di-O-acetyl-l-tartaric anhydride (DATAN), dichloromethane (DCM) and LC/MS grade ammonium acetate were obtained from Sigma (Sigma-Aldrich, St. Louis, MO, USA) for final method optimization. Previously, D-2HG and L-2HG disodium salt was obtained from Toronto Research Chemicals (Toronto, Ontario, Canada). LC/MS grade water, methanol, acetic acid and formic acid were obtained from Covachem (Loves Park, IL, USA). UHPLC/MS Optima acetonitrile was obtained from Fisher Scientific (Hanover Park, IL, USA).

### Sample Preparation for Total D-2-Hydroxyglutarate /L-2-Hydroxyglutarate

The extraction procedure was based on a specimen quantity of ∼3 million cells, ∼16 mg tissue and 200 µL media.

### Nonadherent cell spheroids preparation and collection

Media for specific human-derived stem-like spheroids (BT142, TS603, GSC923) was prepared using 500 mL DMEM: F12 media (Gibco Laboratories, Gaithersburg, MD, USA) with the following additives: 5 mL Penicillin/Streptomycin 100X (Gibco); 5 mL N2 growth supplement 100X (Gibco); 1 mM Heparin Sulfate (Sigma) and 100 µL Epidermal growth factor (EGF) and 100 µL fibroblast growth factors (FGF) (Gibco)

The human derived NCH612 glioma cell media was prepared using 500 mL DMEM: F12/Glutamax media (Gibco) with the following additives: 5 mL Penicillin/Streptomycin 100X; 10 mL B27 supplement 100X (Gibco) and 100 µL EGF and FGF. Following the addition of additives, the media was filter-sterilized and store at 4°C.

Nonadherent cell spheroids were grown in triplicate to a minimum density of 3 million cells per flask. Each sample flask received 12 mL of media and allowed to grow for 72h at 37°C. Samples were transferred to 15-mL pre-sterilized conical tubes and centrifuged at 400 x g for 5 min at room temperature. Then, media from supernatant was collected. Cell pellets were washed with 600 µL PBS and centrifuged at 400 rpm for 5 min at room temperature. Supernatant was discarded. Pellets were snap-frozen on dry ice and stored at -80 °C to ensure complete quenching of all metabolic activity and degradation until extraction.

### Adherent glioma cell preparation and collection

Media for adherent human-derived U251 glioma cells was prepared using 500 mL DMEM (Gibco) with the following additives: 5 mL Penicillin/Streptomycin 100X, 50 µL of 10 mg/mL Puromycin (Gibco); equivalent to 1 µg/mL in media, 1 mM Heparin Sulfate and 50 mL FBS (Gemini Bio-products BenchMark). Following the addition of additives, the media was filter-sterilized and stored at 4°C.

Adherent cells were grown in triplicate to a confluence ≥ 90% in a medium-sized TC-plate. An addition plate should be prepared to approximate cell count and ensure the minimum density of 3.0 million is achieved. Prior to collection, media was collected. Each plate was administered 1.0 mL 0.05% Trypsin and incubated for 1 min. Pipetting was conducted to dissociate adherent cells from plate surfaces. The homogenous cell media was transferred to sterile conical tubes and centrifuged at 400 x g for 5 min at room temperature. Supernatant was discarded and pellet was washed with 1.0 mL PBS. Note: we compared this collection approach to the use of water followed by freezing and scraping cells which did not reveal a significant difference or fold-change in the abundance of D-2HG/L-2HG detected for either preparation (Supplementary Figure S8). Cells were centrifuged 400 x g for 3 min at room temperature. Supernatant was completely aspirated and discarded. The remaining pellet was placed on dry ice to snap-freeze cells and store at -80°C until extraction.

### Media preparation and collection

The all media was aspirated from each cell sample using serological tube to record total volume. 10-mL were transferred to pre-sterilized 15-mL conical tube and placed on dry ice to snap freeze and stored at -80°C. Prior to extraction, media samples were thawed at 4°C and vortexed until homogenized. A 200 µL aliquot was transferred to 1.7-mL microcentrifuge tube and 200 µL of MilliQ pure water was added. A 200 µL aliquot was transferred to 1.7-mL microcentrifuge tube and 200 µL of MilliQ pure water was added to each media sample.

### Tissue preparation and collection

Brain tissue was collected on dry ice to preserve and maintain a solid consistency. A segment of ∼15±5 mg was collected. Each sample measured using an analytical scale. Each sample was placed on dry-ice to snap-freeze and store at -80 °C until extraction.

### Metabolite extraction

#### Sample sonication for cells and tissue

Prior to sonication, tissue and cells samples were administered 400 µL ice-chilled MilliQ water and lysed via sonication by Misonix XL-2000 Ultra-liquid processor (Misonix Inc., Farmingdale, NY, USA) at 40 amps for 30 s. Following sonication, an aliquot (5 % total volume) of each cell lysate was collected for Bradford protein assay to later normalize target analyte concentration to nmol/mg protein.

### Internal standard

Prior to extraction, each remaining 380 µL sample lysate was administered 100 µL of 70% MeOH (aq) internal standard solution—composed of 9.8 nmol/mL U-13C_5_-D-2-Hydroxyglutarate and 9.8 nmol/mL U-13C_5_-L-2-Hydroxyglutarate—chilled at -20°C. Each sample was vortexed on BenchMixer (Benchmark Scientific, Edison, NJ, USA) at mid-speed (6) for 15 s and incubated in ice for 20 min on rotating mixer at mid-speed.

## Conclusions

Our proposed LC/MS methodology detected differentiated and quantified both stereoisomers across all available IDH*^wt^* and IDH*^mut^* cell lines with a count ranging from 3 million cells per sample. Overall, complete reliance on alternate application presents noteworthy limitations when compared to high-throughput performance of LC/MS workflows in conducting 2HG stereoisomer differentiation, biomarker quantification, global metabolome quantitation, comparative analysis, pathway analysis along with novel biomarker and drug-target discovery.

In addition to improvements in resolution for D-/L-enantiomers of 2HG, the proposed DATAN derivatization with LC/MS methodology herein, also facilitated the synchronized quantitation and comparative analysis of other polar biomarkers (i.e. organic acids, amino acids and central carbon intermediates) that would not only be omitted for rapid-result kits given the limited of kits capacity to only quantifying a single biomarker.

D-2-hydroxyglutarate Assay Kit fluorometric and colorimetric applications exists to facilitate benchtop workflows and utilize the specificity of house-keeping enzyme, D-2HG dehydrogenase to indirectly measure D-2HG concentration via its reduction to oxoglutarate and NADH [59]. In comparison to LC/MS-based workflows, these assay kits provide a viable assessment of variability in D-2HG accumulation between specimens, however they are not able to quantify or compare L-2HG concentrations individually or in tandem with D-2HG (D-2-Hydroxyglutarate Assay Kit, Abcam 2017). Therefore, the above method describes a principal advantage of derivatization and follow-on analytical LC/MS methodologies.

While conventional imaging modalities such as positron emission tomography or magnetic resonance spectroscopy have the advantage of non-invasive detection of total 2HG, they do not typically present the necessary resolution to differentiate the D- and L-enantiomers, therefore limiting information can be obtained from the role of each enantiomer to the downstream biological effects [1,11,60–62]. Therefore, the LC/MS methodology proposed by our study fills in this gap.

## Author Contributions

ML and TD have conceived, designed, performed the experiments, and analyzed the data; and conceptualized the study. TD has executed and validated the experiments. TD and ML wrote the paper. ML supervised the study.

## Acknowledgments

We would like to thank Dr. Chun Zhang Yang (Neuro-Oncology Branch, NCI/CCR/NIH) for providing us the U251-WT, U215-R132H and U251-R132C constructs. We also thank Dr. Timothy Chan for providing us with TS603 cell line.

## Ethical Statement

The GSC923 cell line was obtained in-house from the Neuro-Oncology Branch, NCI/CCR/NIH. BT142 was purchased from ATCC. TS603 cell line was obtained from Dr. Timothy Chan, (Memorial Sloan Kettering Cancer Center, New York). U251-WT, U215-R132H and U251-R132C constructs were obtained from Dr. ChunZhang Yang from Neuro-Oncology Branch, NCI/CCR/NIH. Intracranial orthotopic mouse models with an IDH1*^mut^* glioma cell line were established according to an approved animal study proposal NOB-008 by NCI-Animal Use and Care Committee (ACUC).

## References

1. Choi, C.; Ganji, S.K.; DeBerardinis, R.J.; Hatanpaa, K.J.; Rakheja, D.; Kovacs, Z.; Yang, X.L.; Mashimo, T.; Raisanen, J.M.; Marin-Valencia, I., et al. 2-hydroxyglutarate detection by magnetic resonance spectroscopy in IDH-mutated patients with gliomas. Nature medicine 2012, 18, 624–629, doi:10.1038/nm.2682.

2. Dang, L.; White, D.W.; Gross, S.; Bennett, B.D.; Bittinger, M.A.; Driggers, E.M.; Fantin, V.R.; Jang, H.G.; Jin, S.; Keenan, M.C., et al. Cancer-associated IDH1 mutations produce 2-hydroxyglutarate. Nature 2009, 462, 739–744, doi:10.1038/nature08617.

3. Gross, S.; Cairns, R.A.; Minden, M.D.; Driggers, E.M.; Bittinger, M.A.; Jang, H.G.; Sasaki, M.; Jin, S.; Schenkein, D.P.; Su, S.M., et al. Cancer-associated metabolite 2-hydroxyglutarate accumulates in acute myelogenous leukemia with isocitrate dehydrogenase 1 and 2 mutations. The Journal of experimental medicine 2010, 207, 339–344, doi:10.1084/jem.20092506.

4. Amary, M.F.; Bacsi, K.; Maggiani, F.; Damato, S.; Halai, D.; Berisha, F.; Pollock, R.; O’Donnell, P.; Grigoriadis, A.; Diss, T., et al. IDH1 and IDH2 mutations are frequent events in central chondrosarcoma and central and periosteal chondromas but not in other mesenchymal tumours. The Journal of pathology 2011, 224, 334–343, doi:10.1002/path.2913.

5. Kipp, B.R.; Voss, J.S.; Kerr, S.E.; Barr Fritcher, E.G.; Graham, R.P.; Zhang, L.; Highsmith, W.E.; Zhang, J.; Roberts, L.R.; Gores, G.J., et al. Isocitrate dehydrogenase 1 and 2 mutations in cholangiocarcinoma. Human pathology 2012, 43, 1552–1558, doi:10.1016/j.humpath.2011.12.007.

6. Intlekofer, A.M.; Wang, B.; Liu, H.; Shah, H.; Carmona-Fontaine, C.; Rustenburg, A.S.; Salah, S.; Gunner, M.R.; Chodera, J.D.; Cross, J.R., et al. L-2-Hydroxyglutarate production arises from noncanonical enzyme function at acidic pH. Nature chemical biology 2017, 13, 494–500, doi:10.1038/nchembio.2307.

7. Nadtochiy, S.M.; Schafer, X.; Fu, D.; Nehrke, K.; Munger, J.; Brookes, P.S. Acidic pH Is a Metabolic Switch for 2-Hydroxyglutarate Generation and Signaling. The Journal of biological chemistry 2016, 291, 20188–20197, doi:10.1074/jbc.M116.738799.

8. Watanabe, H.; Yamaguchi, S.; Saiki, K.; Shimizu, N.; Fukao, T.; Kondo, N.; Orii, T. Identification of the D-enantiomer of 2-hydroxyglutaric acid in glutaric aciduria type II. Clinica chimica acta; international journal of clinical chemistry 1995, 238, 115–124, doi:10.1016/0009-8981(95)06074-n.

9. Kranendijk, M.; Struys, E.A.; Salomons, G.S.; Van der Knaap, M.S.; Jakobs, C. Progress in understanding 2-hydroxyglutaric acidurias. Journal of inherited metabolic disease 2012, 35, 571–587, doi:10.1007/s10545-012-9462-5.

10. Moroni, I.; Bugiani, M.; D’Incerti, L.; Maccagnano, C.; Rimoldi, M.; Bissola, L.; Pollo, B.; Finocchiaro, G.; Uziel, G. L-2-hydroxyglutaric aciduria and brain malignant tumors: a predisposing condition? Neurology 2004, 62, 1882–1884, doi:10.1212/01.wnl.0000125335.21381.87.

11. Andronesi, O.C.; Loebel, F.; Bogner, W.; Marjanska, M.; Vander Heiden, M.G.; Iafrate, A.J.; Dietrich, J.; Batchelor, T.T.; Gerstner, E.R.; Kaelin, W.G., et al. Treatment Response Assessment in IDH-Mutant Glioma Patients by Noninvasive 3D Functional Spectroscopic Mapping of 2-Hydroxyglutarate. Clinical cancer research : an official journal of the American Association for Cancer Research 2016, 22, 1632–1641, doi:10.1158/1078-0432.Ccr-15-0656.

12. Koivunen, P.; Lee, S.; Duncan, C.G.; Lopez, G.; Lu, G.; Ramkissoon, S.; Losman, J.A.; Joensuu, P.; Bergmann, U.; Gross, S., et al. Transformation by the (R)-enantiomer of 2-hydroxyglutarate linked to EGLN activation. Nature 2012, 483, 484–488, doi:10.1038/nature10898.

13. Bottcher, M.; Renner, K.; Berger, R.; Mentz, K.; Thomas, S.; Cardenas-Conejo, Z.E.; Dettmer, K.; Oefner, P.J.; Mackensen, A.; Kreutz, M., et al. D-2-hydroxyglutarate interferes with HIF-1alpha stability skewing T-cell metabolism towards oxidative phosphorylation and impairing Th17 polarization. Oncoimmunology 2018, 7, e1445454, doi:10.1080/2162402x.2018.1445454.

14. Ye, D.; Guan, K.L.; Xiong, Y. Metabolism, Activity, and Targeting of D- and L-2-Hydroxyglutarates. Trends in cancer 2018, 4, 151–165, doi:10.1016/j.trecan.2017.12.005.

15. Tyrakis, P.A.; Palazon, A.; Macias, D.; Lee, K.L.; Phan, A.T.; Velica, P.; You, J.; Chia, G.S.; Sim, J.; Doedens, A., et al. S-2-hydroxyglutarate regulates CD8(+) T-lymphocyte fate. Nature 2016, 540, 236–241, doi:10.1038/nature20165.

16. Wang, P.; Wu, J.; Ma, S.; Zhang, L.; Yao, J.; Hoadley, K.A.; Wilkerson, M.D.; Perou, C.M.; Guan, K.L.; Ye, D., et al. Oncometabolite D-2-Hydroxyglutarate Inhibits ALKBH DNA Repair Enzymes and Sensitizes IDH Mutant Cells to Alkylating Agents. Cell reports 2015, 13, 2353–2361, doi:10.1016/j.celrep.2015.11.029.

17. Tran, T.Q.; Ishak Gabra, M.B.; Lowman, X.H.; Yang, Y.; Reid, M.A.; Pan, M.; O’Connor, T.R.; Kong, M. Glutamine deficiency induces DNA alkylation damage and sensitizes cancer cells to alkylating agents through inhibition of ALKBH enzymes. PLoS biology 2017, 15, e2002810, doi:10.1371/journal.pbio.2002810.

18. Li, Z.; Weng, H.; Su, R.; Weng, X.; Zuo, Z.; Li, C.; Huang, H.; Nachtergaele, S.; Dong, L.; Hu, C., et al. FTO Plays an Oncogenic Role in Acute Myeloid Leukemia as a N(6)-Methyladenosine RNA Demethylase. Cancer cell 2017, 31, 127–141, doi:10.1016/j.ccell.2016.11.017.

19. Elkashef, S.M.; Lin, A.-P.; Myers, J.; Sill, H.; Jiang, D.; Dahia, P.L.M.; Aguiar, R.C.T. IDH Mutation, Competitive Inhibition of FTO, and RNA Methylation. Cancer cell 2017, 31, 619–620, 10.1016/j.ccell.2017.04.001.

20. Kohanbash, G.; Carrera, D.A.; Shrivastav, S.; Ahn, B.J.; Jahan, N.; Mazor, T.; Chheda, Z.S.; Downey, K.M.; Watchmaker, P.B.; Beppler, C., et al. Isocitrate dehydrogenase mutations suppress STAT1 and CD8+ T cell accumulation in gliomas. The Journal of clinical investigation 2017, 127, 1425–1437, doi:10.1172/jci90644.

21. Su, R.; Dong, L.; Li, C.; Nachtergaele, S.; Wunderlich, M.; Qing, Y.; Deng, X.; Wang, Y.; Weng, X.; Hu, C., et al. R-2HG Exhibits Anti-tumor Activity by Targeting FTO/m(6)A/MYC/CEBPA Signaling. Cell 2018, 172, 90–105.e123, doi:10.1016/j.cell.2017.11.031.

22. Carbonneau, M.; M. Gagné, L.; Lalonde, M.-E.; Germain, M.-A.; Motorina, A.; Guiot, M.-C.; Secco, B.; Vincent, E.E.; Tumber, A.; Hulea, L., et al. The oncometabolite 2-hydroxyglutarate activates the mTOR signalling pathway. Nature Communications 2016, 7, 12700, doi:10.1038/ncomms12700.

23. Chan, S.M.; Thomas, D.; Corces-Zimmerman, M.R.; Xavy, S.; Rastogi, S.; Hong, W.J.; Zhao, F.; Medeiros, B.C.; Tyvoll, D.A.; Majeti, R. Isocitrate dehydrogenase 1 and 2 mutations induce BCL-2 dependence in acute myeloid leukemia. Nature medicine 2015, 21, 178–184, doi:10.1038/nm.3788.

24. Matsunaga, H.; Futakuchi-Tsuchida, A.; Takahashi, M.; Ishikawa, T.; Tsuji, M.; Ando, O. IDH1 and IDH2 have critical roles in 2-hydroxyglutarate production in D-2-hydroxyglutarate dehydrogenase depleted cells. Biochemical and biophysical research communications 2012, 423, 553–556, doi:10.1016/j.bbrc.2012.06.002.

25. Tyrakis, P.A.; Palazon, A.; Macias, D.; Lee, K.L.; Phan, A.T.; Veliça, P.; You, J.; Chia, G.S.; Sim, J.; Doedens, A., et al. S-2-hydroxyglutarate regulates CD8+ T-lymphocyte fate. Nature 2016, 540, 236–241, doi:10.1038/nature20165.

26. Chowdhury, R.; Yeoh, K.K.; Tian, Y.M.; Hillringhaus, L.; Bagg, E.A.; Rose, N.R.; Leung, I.K.; Li, X.S.; Woon, E.C.; Yang, M., et al. The oncometabolite 2-hydroxyglutarate inhibits histone lysine demethylases. EMBO reports 2011, 12, 463–469, doi:10.1038/embor.2011.43.

27. Ma, S.; Sun, R.; Jiang, B.; Gao, J.; Deng, W.; Liu, P.; He, R.; Cui, J.; Ji, M.; Yi, W., et al. L2hgdh Deficiency Accumulates l-2-Hydroxyglutarate with Progressive Leukoencephalopathy and Neurodegeneration. Molecular and cellular biology 2017, 37, doi:10.1128/mcb.00492-16.

28. Struys, E.A.; Jansen, E.E.; Verhoeven, N.M.; Jakobs, C. Measurement of urinary D- and L-2-hydroxyglutarate enantiomers by stable-isotope-dilution liquid chromatography-tandem mass spectrometry after derivatization with diacetyl-L-tartaric anhydride. Clinical chemistry 2004, 50, 1391–1395, doi:10.1373/clinchem.2004.033399.

29. Poinsignon, V.; Mercier, L.; Nakabayashi, K.; David, M.D.; Lalli, A.; Penard-Lacronique, V.; Quivoron, C.; Saada, V.; De Botton, S.; Broutin, S., et al. Quantitation of isocitrate dehydrogenase (IDH)-induced D and L enantiomers of 2-hydroxyglutaric acid in biological fluids by a fully validated liquid tandem mass spectrometry method, suitable for clinical applications. Journal of chromatography. B, Analytical technologies in the biomedical and life sciences 2016, 1022, 290–297, doi:10.1016/j.jchromb.2016.04.030.

30. Cheng, Q.-Y.; Xiong, J.; Huang, W.; Ma, Q.; Ci, W.; Feng, Y.-Q.; Yuan, B.-F. Sensitive Determination of Onco-metabolites of D- and L-2-hydroxyglutarate Enantiomers by Chiral Derivatization Combined with Liquid Chromatography/Mass Spectrometry Analysis. Scientific reports 2015, 5, 15217, doi:10.1038/srep15217.

31. Oldham, W.M.; Loscalzo, J. Quantification of 2-Hydroxyglutarate Enantiomers by Liquid Chromatography-mass Spectrometry. Bio-protocol 2016, 6, e1908, doi:10.21769/BioProtoc.1908.

32. Schreiber , A. Advantages of Using Triple Quadrupole over Single Quadrupole Mass Spectrometry to Quantify and Identify the Presence of Pesticides in Water and Soil Samples. AB Sciex 2010, 0701310-01, 1–6.

33. Xia, Y.Q.; Lau, J.; Olah, T.; Jemal, M. Targeted quantitative bioanalysis in plasma using liquid chromatography/high-resolution accurate mass spectrometry: an evaluation of global selectivity as a function of mass resolving power and extraction window, with comparison of centroid and profile modes. Rapid communications in mass spectrometry : RCM 2011, 25, 2863–2878, doi:10.1002/rcm.5178.

34. Wishart, D.S.; Feunang, Y.D.; Marcu, A.; Guo, A.C.; Liang, K.; Vazquez-Fresno, R.; Sajed, T.; Johnson, D.; Li, C.; Karu, N., et al. HMDB 4.0: the human metabolome database for 2018. Nucleic acids research 2018, 46, D608–d617, doi:10.1093/nar/gkx1089.

35. Chong, J.; Wishart, D.S.; Xia, J. Using MetaboAnalyst 4.0 for Comprehensive and Integrative Metabolomics Data Analysis. Current Protocols in Bioinformatics 2019, 68, e86, doi:10.1002/cpbi.86.

36. Xia, Y.-Q.; Lau, J.; Olah, T.; Jemal, M. Targeted quantitative bioanalysis in plasma using liquid chromatography/high-resolution accurate mass spectrometry: an evaluation of global selectivity as a function of mass resolving power and extraction window, with comparison of centroid and profile modes. Rapid Communications in Mass Spectrometry 2011, 25, 2863–2878, doi:10.1002/rcm.5178.

37. The Theory of HPLC Chromatographic Parameters. academy, C., Ed.

38. Kranendijk, M.; Salomons, G.S.; Gibson, K.M.; Van Schaftingen, E.; Jakobs, C.; Struys, E.A. A lymphoblast model for IDH2 gain-of-function activity in d-2-hydroxyglutaric aciduria type II: Novel avenues for biochemical and therapeutic studies. Biochimica et Biophysica Acta (BBA) - Molecular Basis of Disease 2011, 1812, 1380–1384, 10.1016/j.bbadis.2011.08.006.

39. Jones, P.M.; Boriack, R.; Struys, E.A.; Rakheja, D. Measurement of Oncometabolites D-2-Hydroxyglutaric Acid and L-2-Hydroxyglutaric Acid. Methods in molecular biology (Clifton, N.J.) 2017, 1633, 219–234, doi:10.1007/978-1-4939-7142-8_14.

40. Kalinina, J.; Ahn, J.; Devi, N.S.; Wang, L.; Li, Y.; Olson, J.J.; Glantz, M.; Smith, T.; Kim, E.L.; Giese, A., et al. Selective Detection of the D-enantiomer of 2-Hydroxyglutarate in the CSF of Glioma Patients with Mutated Isocitrate Dehydrogenase. Clinical cancer research : an official journal of the American Association for Cancer Research 2016, 22, 6256–6265, doi:10.1158/1078-0432.Ccr-15-2965.

41. Larion, M.; Dowdy, T.; Ruiz-Rodado, V.; Meyer, M.W.; Song, H.; Zhang, W.; Davis, D.; Gilbert, M.R.; Lita, A. Detection of Metabolic Changes Induced via Drug Treatments in Live Cancer Cells and Tissue Using Raman Imaging Microscopy. Biosensors 2018, 9, doi:10.3390/bios9010005.

42. Huang, J.; Zhao, Q.; Mooney, S.M.; Lee, F.S. Sequence determinants in hypoxia-inducible factor-1alpha for hydroxylation by the prolyl hydroxylases PHD1, PHD2, and PHD3. The Journal of biological chemistry 2002, 277, 39792–39800, doi:10.1074/jbc.M206955200.

43. Tateishi, K.; Wakimoto, H.; Iafrate, A.J.; Tanaka, S.; Loebel, F.; Lelic, N.; Wiederschain, D.; Bedel, O.; Deng, G.; Zhang, B., et al. Extreme Vulnerability of IDH1 Mutant Cancers to NAD+ Depletion. Cancer cell 2015, 28, 773–784, doi:10.1016/j.ccell.2015.11.006.

44. Zhou, Z.; Ibekwe, E.; Chornenkyy, Y. Metabolic Alterations in Cancer Cells and the Emerging Role of Oncometabolites as Drivers of Neoplastic Change. Antioxidants (Basel, Switzerland) 2018, 7, doi:10.3390/antiox7010016.

45. Vander Heiden, M.G.; DeBerardinis, R.J. Understanding the Intersections between Metabolism and Cancer Biology. Cell 2017, 168, 657–669, doi:10.1016/j.cell.2016.12.039.

46. Nowicki, S.; Gottlieb, E. Oncometabolites: tailoring our genes. The FEBS journal 2015, 282, 2796–2805, doi:10.1111/febs.13295.

47. Pavlova, N.N.; Thompson, C.B. The Emerging Hallmarks of Cancer Metabolism. Cell metabolism 2016, 23, 27–47, doi:10.1016/j.cmet.2015.12.006.

48. Ohka, F.; Ito, M.; Ranjit, M.; Senga, T.; Motomura, A.; Motomura, K.; Saito, K.; Kato, K.; Kato, Y.; Wakabayashi, T., et al. Quantitative metabolome analysis profiles activation of glutaminolysis in glioma with IDH1 mutation. Tumour biology : the journal of the International Society for Oncodevelopmental Biology and Medicine 2014, 35, 5911–5920, doi:10.1007/s13277-014-1784-5.

49. Victor, R.R.; Malta, T.M.; Seki, T.; Lita, A.; Dowdy, T.; Celiku, O.; Cavazos-Saldana, A.; Li, A.; Liu, Y.; Han, S., et al. Metabolic Reprogramming Associated with Aggressiveness Occurs in the G-CIMP-High Molecular Subtypes of IDH1mut Lower Grade Gliomas. Neuro-oncology 2019, 10.1093/neuonc/noz207, doi:10.1093/neuonc/noz207.

50. Losman, J.A.; Kaelin, W.G., Jr. What a difference a hydroxyl makes: mutant IDH, (R)-2-hydroxyglutarate, and cancer. Genes & development 2013, 27, 836–852, doi:10.1101/gad.217406.113.

51. Rogatzki, M.J.; Ferguson, B.S.; Goodwin, M.L.; Gladden, L.B. Lactate is always the end product of glycolysis. Frontiers in neuroscience 2015, 9, 22, doi:10.3389/fnins.2015.00022.

52. Jeoung, N.H. Pyruvate Dehydrogenase Kinases: Therapeutic Targets for Diabetes and Cancers. Diabetes & metabolism journal 2015, 39, 188–197, doi:10.4093/dmj.2015.39.3.188.

53. De Saedeleer, C.J.; Copetti, T.; Porporato, P.E.; Verrax, J.; Feron, O.; Sonveaux, P. Lactate activates HIF-1 in oxidative but not in Warburg-phenotype human tumor cells. PloS one 2012, 7, e46571, doi:10.1371/journal.pone.0046571.

54. Mazor, T.; Chesnelong, C.; Pankov, A.; Jalbert, L.E.; Hong, C.; Hayes, J.; Smirnov, I.V.; Marshall, R.; Souza, C.F.; Shen, Y., et al. Clonal expansion and epigenetic reprogramming following deletion or amplification of mutant IDH1. Proceedings of the National Academy of Sciences of the United States of America 2017, 114, 10743–10748, doi:10.1073/pnas.1708914114.

55. Luchman, H.A.; Chesnelong, C.; Cairncross, J.G.; Weiss, S. Spontaneous loss of heterozygosity leading to homozygous R132H in a patient-derived IDH1 mutant cell line. Neuro-oncology 2013, 15, 979–980, doi:10.1093/neuonc/not064.

56. Fan, J.; Kamphorst, J.J.; Mathew, R.; Chung, M.K.; White, E.; Shlomi, T.; Rabinowitz, J.D. Glutamine-driven oxidative phosphorylation is a major ATP source in transformed mammalian cells in both normoxia and hypoxia. Molecular systems biology 2013, 9, 712, doi:10.1038/msb.2013.65.

57. Rajagopalan, K.N.; Egnatchik, R.A.; Calvaruso, M.A.; Wasti, A.T.; Padanad, M.S.; Boroughs, L.K.; Ko, B.; Hensley, C.T.; Acar, M.; Hu, Z., et al. Metabolic plasticity maintains proliferation in pyruvate dehydrogenase deficient cells. Cancer & metabolism 2015, 3, 7, doi:10.1186/s40170-015-0134-4.

58. Elhammali, A.; Ippolito, J.E.; Collins, L.; Crowley, J.; Marasa, J.; Piwnica-Worms, D. A high-throughput fluorimetric assay for 2-hydroxyglutarate identifies Zaprinast as a glutaminase inhibitor. Cancer discovery 2014, 4, 828–839, doi:10.1158/2159-8290.Cd-13-0572.

59. Chua, Y.L.; Dufour, E.; Dassa, E.P.; Rustin, P.; Jacobs, H.T.; Taylor, C.T.; Hagen, T. Stabilization of hypoxia-inducible factor-1alpha protein in hypoxia occurs independently of mitochondrial reactive oxygen species production. The Journal of biological chemistry 2010, 285, 31277–31284, doi:10.1074/jbc.M110.158485.

60. Kim, M.M.; Lawrence, T.S.; Cao, Y. Advances in Magnetic Resonance and Positron Emission Tomography Imaging: Assessing Response in the Treatment of Low-Grade Glioma. Seminars in radiation oncology 2015, 25, 172–180, doi:10.1016/j.semradonc.2015.02.003.

61. Elkhaled, A.; Jalbert, L.E.; Phillips, J.J.; Yoshihara, H.A.I.; Parvataneni, R.; Srinivasan, R.; Bourne, G.; Berger, M.S.; Chang, S.M.; Cha, S., et al. Magnetic resonance of 2-hydroxyglutarate in IDH1-mutated low-grade gliomas. Science translational medicine 2012, 4, 116ra115, doi:10.1126/scitranslmed.3002796.

62. Andronesi, O.C.; Arrillaga-Romany, I.C.; Ly, K.I.; Bogner, W.; Ratai, E.M.; Reitz, K.; Iafrate, A.J.; Dietrich, J.; Gerstner, E.R.; Chi, A.S., et al. Pharmacodynamics of mutant-IDH1 inhibitors in glioma patients probed by in vivo 3D MRS imaging of 2-hydroxyglutarate. Nature Communications 2018, 9, 1474, doi:10.1038/s41467-018-03905-6.

