## Supplemental methods and figures for "Resolving Challenges in Detection and Quantification of D-2-hydroxyglutarate and L-2-hydroxyglutarate via LC/MS"

### Metabolite extraction of tissue and cell

Upon extraction, 600 µL chilled (-20 °C) LC/MS grade Methanol (MeOH) was added to each 480 µL sample lysate with added IS. Samples were vortexed with for 30 s and placed on Orbi-blotter mixing rotator (Benchmark Scientific, Edison, NJ, USA) at max speed to incubate in ice-bath for 10 min. Samples were vortexed and administered 300 µL chilled (-20 °C) acetonitrile (ACN) each and returned to ice-bath on rotator for 30 min to precipitate protein.

### Extract collection

All samples remained in ice-bath until final centrifugation and collection. Samples were centrifuged at 14200 x g for 20 min at 4°C. Following centrifugation, 90% of the supernatant volume was transferred to new tube while aspirating slowly to not disrupt protein pellet. Extracts were placed under $N_2$ gas flow on Techne sample concentrator with PTFE-coated needles (Cole-Palmer, Vernon Hills, IL, USA) at 25 °C until completely dry and then stored at -80 °C.

### Chiral derivatization

Extracts with derivatized to resolve D-2HG and L-2HG enantiomers in addition to structural isomers of these analytes via esterification with DATAN.

1. Stock solutions of D-2HG and L-2HG and the $^{13}C_5$-2HG internal standards (IS) were prepared with MilliQ pure $H_2O$ for immediate use of working solutions for standard curve and excess was stored at -20 °C.
2. 9.8 nmol/mL $^{13}C_5$-D-2HG and $^{13}C_5$-L-2HG IS solution was prepared in 80% MeOH (aq)
   - Derivatization reagent was prepared at 50 mg/mL DATAN in 80% acetonitrile and 20 % acetic acid for immediate sample preparation. Only freshly prepared derivatizing DATAN reagent was used (up to 24h).
3. Standard solutions and extracts were reconstituted in 60 µL of DATAN reagent on the day of experiment. (Note: If samples were in freezer prior to derivatization, place them under $N_2$ gas stream for 15 min and ensure that no moisture remains. Water impedes the efficacy of the derivatization process.
4. Standards and extracts were vortexed for 1 min and centrifuged at 22400 x g for 30 s at 4 °C.
5. Then, incubated at 75°C with continuous shaking at 1400 rpm for 32 min on ThermoMixer (Eppendorf, Hauppauge, NY).
6. Eppendorf's caps were tightened and centrifuged at 22400 x g for 30 s at 4°C to cool and pool samples+ (Note: Derivatized standards present clear to faint yellow. Derivatized samples typically present light yellow while saturated samples present yellow-brown hue).
7. Samples were concentrated for ~2h on SpeedVac (Eppendorf, Hauppauge, NY, USA) at 35°C until dryness (Note: A greasy residue will remain in tube. Upon completion, pluck bottom of tube to ensure that residue does not splash. If residue splash, continue to dry for additional 30 min).
8. Dried samples were reconstituted in 100 µL of 100 % $H_2O$ (LC/MS grade), homogenized via vortexing (15 s on the high speed) and centrifuged at 10000 x g for 5 min at 4°C.
9. Samples were transferred to amber glass LC vial with deactivated 250-µL inserts (Agilent Technologies, Wilmington, DE, USA). (Note: Aspirate slowly from top-down to not disrupt white pellet in microcentrifuge tube).

### LC/MS analysis of D-2HG and L-2HG

Superior separation of D- and L- stereoisomers of 2-HG was achieved with Hi-Resolution Agilent 6545 Quadrupole Time-of-Flight (QTOF) Liquid Chromatography Mass Spectrometry with Liquid Chromatography (LC/MS) system using the novel gradient and application of Zorbax Eclipse RRHD 2.1 x 100 mm, 1.8µm column (Agilent Technologies, Wilmington, DE, USA).

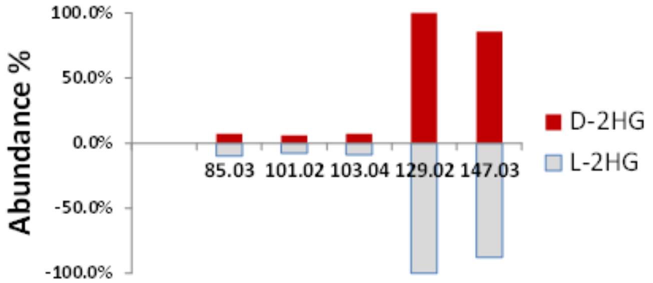


Figure 12. Comparison of LCMS/MS collision-induced dissociation (CID) fragmentation pattern for D-2HG/L-2HG at collision energy (CE) of 2 V following derivatization and LC/MS analysis while applying the condition described in Methods section.

Prior to quantitative analysis, targeted tandem MS (MS/MS) was applied (Figure 12) to validate the fragmentation of D-2HG and L-2HG analogs which are the major fragment the 2HG-DATAN esters (m/z 363.057) as determined in previous publications (28). Based on enantioselective detection of their respective precursor ions (m/z 147.030), both analytes presented a major transition ion (m/z 129.018) which was referenced as a qualifier for MS/MS analysis in negative electrospray ionization mode (ESI-). The concentrations for D-2HG and L-2HG were normalized to area of internal standard (IS), U-13C-D-2HG or U-13C-L-2HG, respectively. Targeted MS/MS analysis was also applied to validate the detection and retention time of U-13C-2HG analogs of the U13C-2HG-DATAN standards based on detection of respective precursor ion (m/z 152.047) and transition ion (m/z 134.036) as a qualifier for ESI- analysis. Under the LC/MS experimental conditions, D-2HG and U13C-D-2HG (IS) co-eluted at 6 min; while L-2HG and U13C-L2HG (IS) co-eluted at 5 min. Targeted compound detection and peak integration was achieved with Masshunter Profinder B.10.0. All solvents and additives used were LC/MS grade, unless indicated as otherwise.

- Instrumentation:
  - 6545 Quadrupole Time of Flight Mass Spectrometer coupled with 1290 Infinity II Ultra-High-Pressure Liquid Chromatography Unit (Agilent Technologies, Santa Clara, CA, USA).
- LC conditions
  - Column: Zorbax Eclipse 2.1 x 100 mm 1.8μm and Eclipse 2.1 mm 1.8μm guard C18 columns (Agilent Technologies, Wilmington, DE, USA)
  - Column temperature: 40 °C
  - Mobile phase A (LC/MS grade): 2 mM Ammonium acetate in 100% H2O (Fisher Optima) pH 3.5 titrated with formic acid (LC/MS grade, Covachem)
  - Mobile phase B (LC/MS grade): 95% MeOH (Covachem)/ 5% ACN (Fisher UHPLC Optima)
  - Flow rate: 0.200 mL/min
  - Gradient: Initial 3% B hold for 6 min; ramp to 5% B min over 1 min; ramp to 99% B over 5 min; hold for 1 min; equilibrate at initial conditions for 2.5 min.
  - • Injection 6 μL
- MS conditions:
  - 3.8 spectra/s
  - VCap: 2600 V
  - Nozzle: 1000 V
  - Fragmentor: 80 V
  - Skimmer: 37 V
  - Drying cap gas: flow 8 L/min at 300 °C
  - Sheath gas: flow 10 L/min at 250 °C
  - Nebulizer 45 psig
  - Collision energy: 0 V

**Quantitative Analysis**

Standard curves and sample calibrations for 2HG were calculated by plotting the ratios for peak area for D-2HG/L-2HG standards (0-104 nmol/mL) separately over respective internal standards and fitting with normal, unweighted linear regression in Excel. Graphs were generated in GraphPad Prism v7.05. Multivariate biostatistical analysis was performed using Metabolanalyst 4.0 application with Pearson-Ward correlation analysis [35]. Statistical analysis conducted by applying non-parametric, independent samples t-tests for each metabolite area abundance adjusted to internal standard response per sample. Relative abundances were generated upon correction of adjusted abundances by sample protein content (mg) measured via Bradford protein quantification.

**Supplementary Figure S1**:

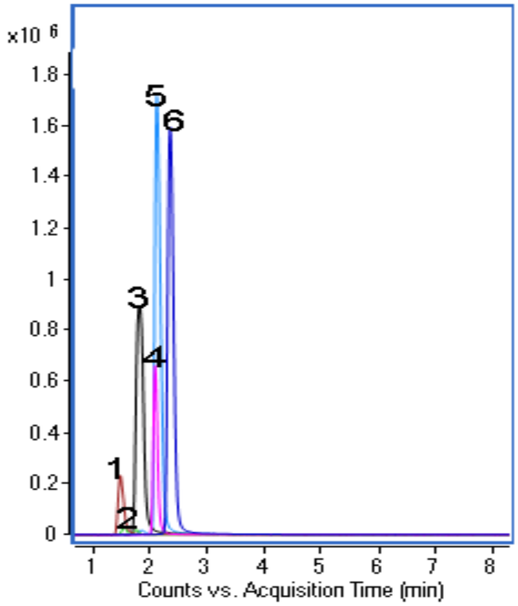


Supplementary Figure S1. XICs for non-derivatized standards for enantiomers D-2HG and L-2HG (3) and related endogenous structural isomer: 1) Ribonolactone; 2) Xylonolactone; 4) 3-Hydroxyglutarate; 5) 3-Methylmalate; 6) Citramalate (2-Methylmalate). Each non-derivatized standard prepared at 26.0 nmol/mL was injected and analyzed under the same LC/MS conditions as the optimized D-2HG and L-2HG quantitative analysis method described in Methods section. Standards were prepared at 26 nmol/mL without chiral derivatization. Standards were injected at 6μL as indicated in Methods section.

**Supplementary Figure S2**:

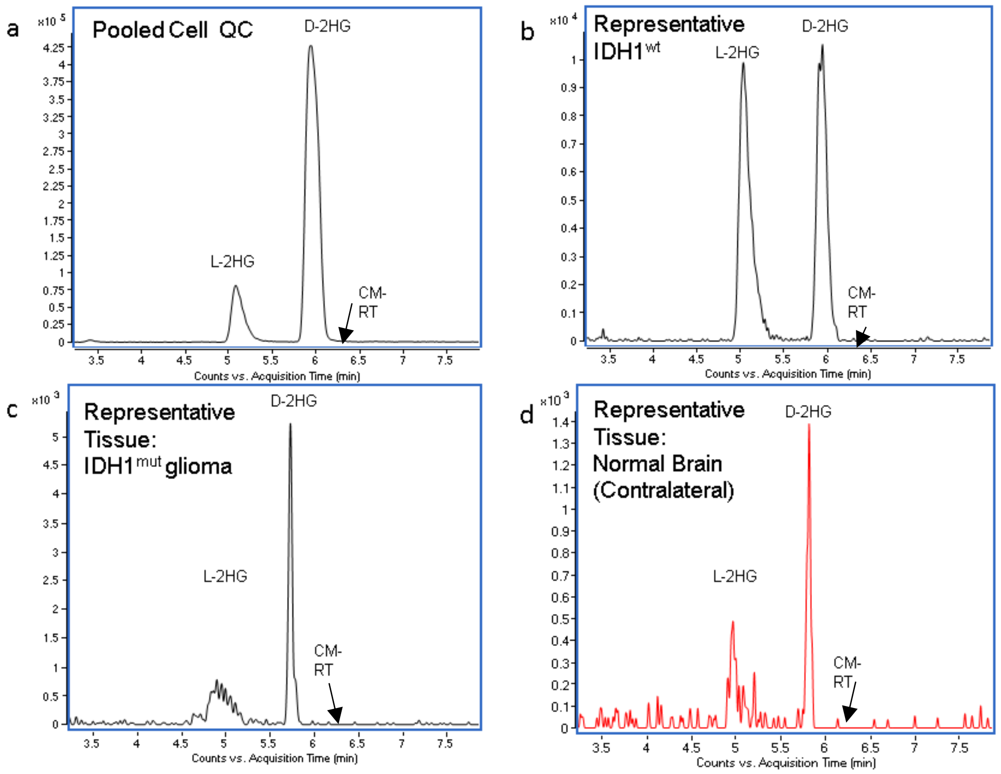


**Supplementary Figure S2**. Extracted ion chromatograms (XICs) a) for L-2HG and D-2HG (black, left to right) in biological samples following sample preparation, chiral derivatization with DATAN and analyzed according to the proposed methodology in Methods section. Qualitative scan of XICs at m/z 147.030 for a) pooled sample QC of both, IDH1 mutant glioma and wild-type GBM cells; b) IDH$^{wt}$ only; c) IDH1$^{mut}$ brain tumor from mice; d) normal brain tissue from mice did not reveal any evident, interfering peak at the validated retention time (~6.3 min) for citramalic acid (CM-RT).

**Supplementary Figure S3:**

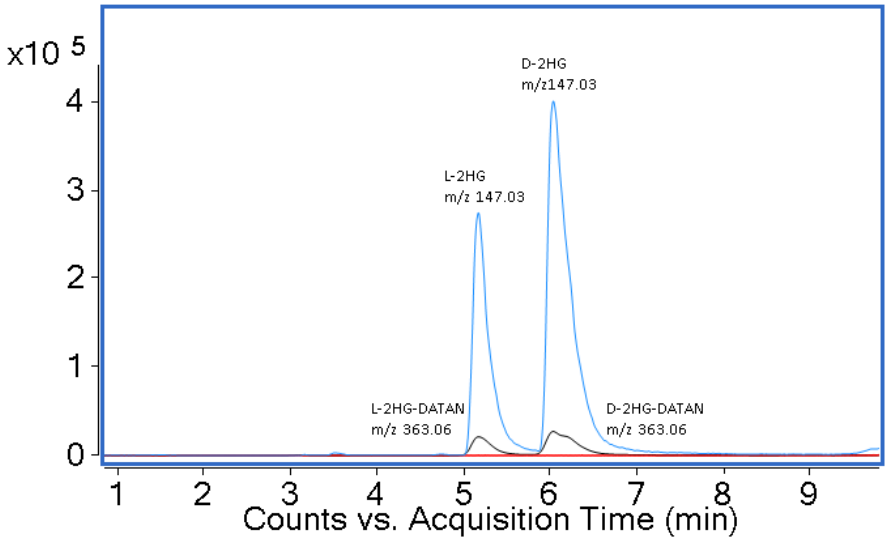


**Supplementary Figure S3.** Carryover following high-concentration injection of racemic 2HG (rac-2HG). An XIC overlay of D-2HG/L-2HG analog (blue), 2HG-DATAN esters (black), as well as the consecutive blank (red) are shown. D-2HG/L-2HG and 2HG-DATAN detected in blank appears to be below limits of detection immediately following high-concentration injection of 73 nmol/mL rac-2HG using TOF-MS analysis (CE = 0V) after chiral derivatization with DATAN and analyzed vis LC/MS on Zorbax

Eclipse C18 column as described in Methods section. Ion abundance is represented in counts per sec (y-axis) and the elution time is represented in minutes (x-axis).

**Supplementary Figure S4:**

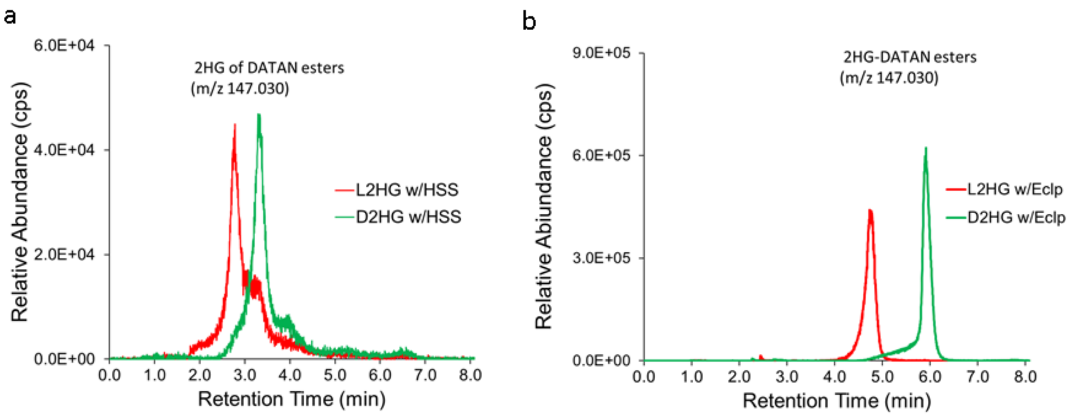


**Supplementary Figure S4**. The chromatograms for acquisition on the a) Xbridge HSS T3 C18 and b) Eclipse Plus RRHD C18 columns illustrated the best separation of D-2HG-DATAN and L-2HG-DATAN for the two column that performed better than the other columns tested using reverse phase gradient as described above (Supplementary Table S2). The ion abundance in counts per second (cps) is indicated along the y-axis and the retention time in minutes (min) along the x-axis. Standards were prepared at 52 nmol/mL and derivatized with DATAN as explained in Methods section. Supplementary Table S3 provides details for the reverse phase gradient with ammonium formate additive that was separate these analytes.

**Supplementary Figure S5:**

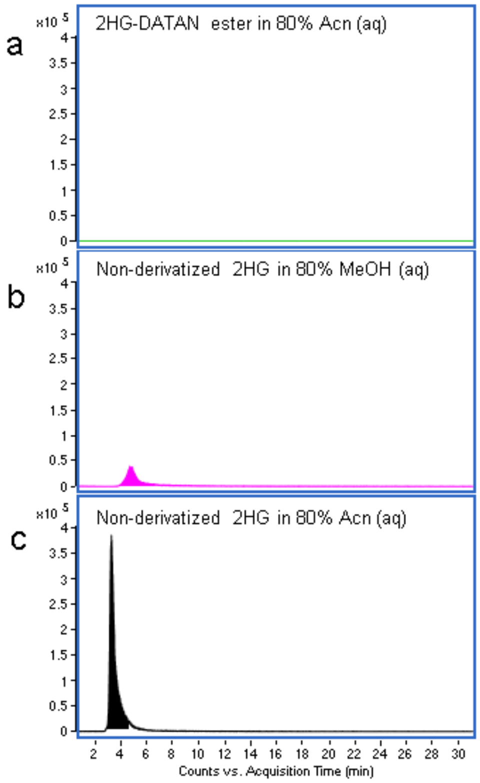

**Supplementary Figures S5.** Sequant Zwitterionic HILIC (HILIC-Z) column was tested to examine XIC (m/z 147.030) for 16 nmol/mL pure D-2HG/L-2HG standards that were derivatized as described in Methods section. a) Main XIC for racemic-2HG derivatized in DATAN and resuspended in 80% ACN which revealed poor retention with abundance counts below detection limits; b) preliminary XIC for non-derivatized 2HG in 80% MeOH diluent; c) preliminary XIC for non-derivatized 2HG in 80% ACN diluent which presented best retention and chromatography under HILIC condition. Each standard was injected (5µL) on Sequant HILIC-Z 2.1x150mm column (Supelco Inc. Bellefonte, PA, USA) over a 37 min gradient using mobile phase composed of A (20 mM FA in $H_20$ pH 3.25) and B (90:10 ACN/20 mM FA in $H_20$ pH 3.25). Analysis was conducted in ESI- mode as described in Methods section. Hilic-Z gradient parameters were as indicated here: 85% B initial; hold 30 min; ramp to 50% B by 33 min; ramp to 25% B by 37 min. The XIC provide abundance in counts per second (y-axis) and elution time in minutes (x-axis).

**Supplementary Figure S6:**

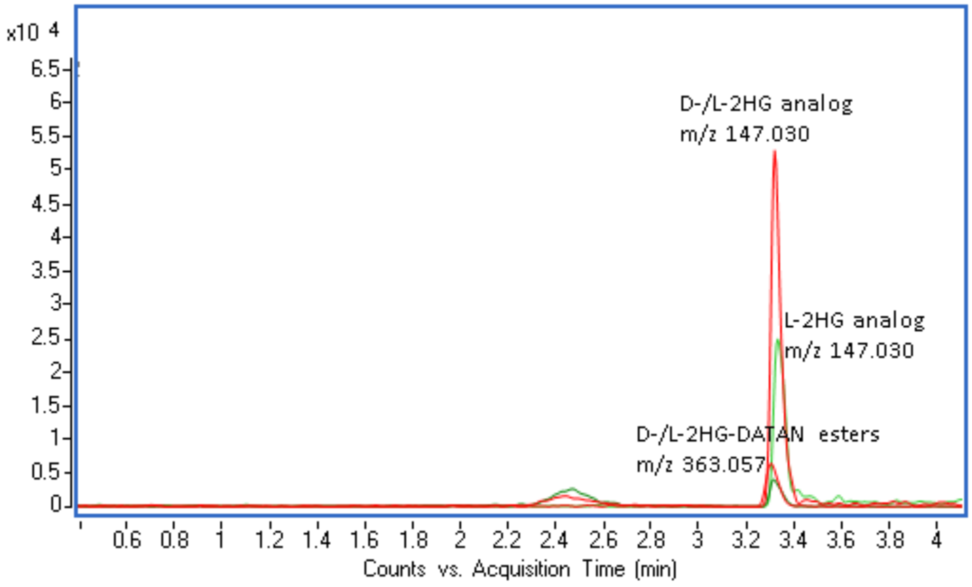


**Supplementary Figures S6.** AdvanceBio Glycan HILIC column experimental XIC overlay for (m/z 147.030) 8 nmol/mL standard D-2HG analog (147.03) along with D-2HG-DATAN ester (m/z 363.057) in green and L-2HG analog (147.030) along with L-2HG-DATAN ester (m/z 363.057) in red that were derivatized as described in Methods section and resuspended in 50% ACN which reveals poor separation while better retention and ion abundance than Sequant HILIC-Z experiment (even at a half concentration). Each standard was injected (5µL) on 10.5 min gradient using mobile phases A (12.0 % ACN in Water with 0.01%v/v 5 mM Medronic acid) and B (90:10 ACN/Water) both phase at 10 mM ammonium acetate; both titrated with ammonium hydroxide and formic acid to pH 6.75. Analysis was conducted in ESI- mode (as described in Methods section). Glycan HILIC gradient parameters were as indicated here: 100% B initial; hold 0.5 min; ramp to 57% B by 2 min; hold 2.5 min; ramp to 56% B by 5 min; ramp to 20 % B by 9.5 min, ramp to 12% B by 10.5 min. The XIC provide abundance in counts per second (y-axis) and elution time in minutes (x-axis).

**Supplementary Figure S7:**

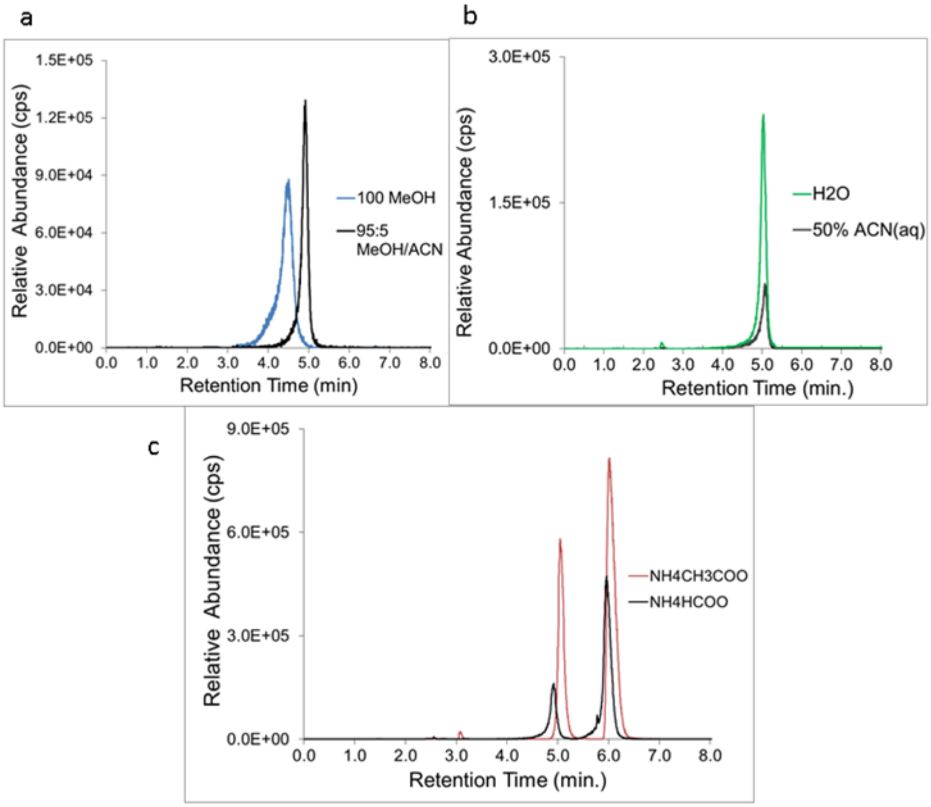


**Supplementary Figure S7.** The chromatograms compare experimental conditions that were surveyed to optimize the resolution and chromatography after the optimal column Eclipse Plus RRHD and gradient were determined as indicated in caption for Supplementary Figure S4 and Table S2. a) The final composition of solvent B (100% MeOH vs. 95:5 MeOH/ACN) was assessed with L-2HG. b) The impact on ionization of L-2HG is illustrated when the diluents composition is completely aqueous at 100% H2O vs. partially aqueous at 50% ACN. c) The use of solvent additives 2 mM ammonium acetate (NH4CH3COO) and 2 mM ammonium formate (NH4HCOO) in mobile phase A was compared to optimize separation, retention and ionization for 2HG enantiomers. Based on data illustrated in the XIC, we adapted solvent B to 95:5 MeOH/ACN and mobile phase A to 2 mM ammonium acetate (pH 3.5) in $H_2O$. These XICs serve as visual representations for the main relevant changes observed from our comparison of chromatographic chemistry factors as detailed in Supplementary Table S2 (above). Unless otherwise indicated, the same reverse phase was used. Standards were prepared at 52 nmol/mL and derivatized as described in Methods section using reverse phase gradient described for Supplementary Table S3. The ion abundance in counts per second (cps) is indicated along the y-axis and the retention time in minutes (min) along the x-axis.

**Supplementary Figure S8:**

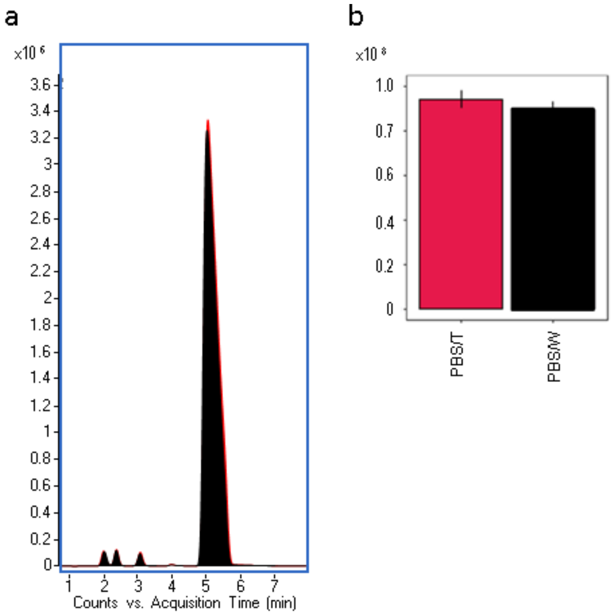


**Supplementary Figure S8.** Ideal Collection Method Comparison for PBS+Trypsin vs PBS+Water. Overlay of extracted ion chromatogram a) for six biological replicate samples of adherent wild-type U251 glioma cells with genetic overexpression of IDH1-R132H mutation that was prepared in two sets of three to facilitate qualitative and statistical comparison of two different collection techniques: the use of PBS with trypsin and no water (red); and the use of PBS follow by wash with water (black) to reduce PBS salt-contamination. The graph in b) shows average peak area for extracted ions at m/z 147.030 for each group as a function of the two different collection methods. PBS/T, cell collection using PBS, trypsin, and no water (n=3); PBS/W, cell collection using PBS and MilliQ water (n=3).

Two separate sample collection approaches were compared as indicated: 1) PBS/T involved the use of Trypsin, PBS, and centrifuge to retain only the pellet in tube; and 2) PBS/W involved PBS wash followed by brief wash with low volume of water (to reduce suppression by salt residues from PBS), snap-freezing on dry-ice and cell scraping from plate (Supplementary Figure S8a-b). The differences in the detection were unremarkably low as (Supplementary Figure S8a). The data in the graph (Supplementary Figure S8b) revealed an insignificant fold-change of 1.05 (Welch's t-test p-value = 0.21) for the detection of D-2HG (m/z 147.030) between the two approaches. Therefore, the use of PBS with or without water did not appear to significantly impact the detection of the target analytes. We adapted the method to incorporate that approach which combines the use of Trypsin, PBS and centrifugation given that it reduces sample preparation time and potential for sample loss.

**Supplementary Table S1.** Precision (intra- and inter-day) for determination of D-2HG and L-2HG in pooled sample QC composed of glioma and GBM cells derivatized with DATAN and analyzed via LC/MS analysis as described in Methods section.

| **Analyte** | **Intra-day 1** CV% (n = 2) | **Intra-day 2** CV% (n = 2) | **Avg Intra-day** CV% (n = 4) | **Inter-day (2 day)** CV% (n = 4) |
|---|---|---|---|---|
| **D-2HG** | 1.11 | 0.01 | 0.56 | 0.70 |
| **L-2HG** | 1.13 | 0.01 | 0.57 | 0.80 |

**Supplementary Table S2**. DATAN derivatization conditions tested to maximize reproducibility, chemical equilibrium and stability. *

| Derivatization Application | Conditions |
|---|---|
| **Derivatizing agent** | DATAN in 1:4 Acetic acid/ Acetonitrile |
| | DATAN in 1:4 Acetic acid/ DCM |
| **DATAN concentration (mM)** | 115, 230 |
| **Volume (µL)** | 50, 60, 100 |

| | |
|---|---|
| **Catalyst** | 2 µM Sodium Lactate (aq), None |
| **Temperature (°C)** | 70, 75 |
| **Incubation and Mixing time (min)** | 32, 60, 90 |
| **Mixing speed (rpm)** | 0, 850, 1400 |
| **Drying application (Temperature)** | $N_2$ gas concentrator (25°C, 35°C) |
| | Speed vac concentrator (25°C, 35°C) |

*Based on these comparisons in Supplementary Table S2, it was determined that complete chiral derivatization and subsequent enantiomeric separation of D-2HG/L-2HG can be achieved added 60 µL of 230 mM DATAN in 1:4 Acetic acid/ DCM without catalyst to each sample and standard; followed by incubated for 32 min at 75ºC while mixing at 1400 rpm with drying time (~2h) with SpeedVac at 35 ºC varying with sample composition.

**Supplementary Table S3**. LC conditions tested and compared to maximize separation, detection and chromatography*chromatography. *

| LC-MS Application | Conditions |
|---|---|
| Aqueous buffers | 2 mM Ammonium formate pH 3.5 |
| | 2 mM Ammonium acetate pH 3.5 |
| | 20 mM Formic acid pH 3.25 |
| | 0.1%v/v Formic acid + 0.01%v/v 5 mM Medronate |
| Organic buffers | 95:5 MeOH/ACN |
| | 95:5 ACN/MeOH |
| | 50:50 ACN/MeOH |
| | 100% MeOH |
| | 90:10 ACN/ 200 mM Formic acid (aq) |
| Columns | Sequant HilicZ, 2.1x150mm, 5µm with HilicZ guard (Supelco) |
| | Discovery HS F5, 2.1x100mm, 3µm (Supelco) |
| | Xselect BEH phenyl C18, 2.1x50mm, 2.5 µm (Waters) |
| | Xselect HSS T3 C18, 2.1x100mm, 2.5 µm (Waters) |
| | AdvanceBio Glycan Map, 2.1x100mm, 2.7 µm (Agilent) |
| | Zorbax EclipsePlus RRHD C18, 2.1x100mm, 1.8 µm (Agilent) |
| Diluents | 99:1 $H_2O$/ Acetic acid |
| | 4:1 ACN/ Acetic acid |
| | 60% MeOH (aq) |
| | 50% ACN (aq) |
| | 80% ACN (aq) |
| | 80% MeOH (aq) |
| | 100% $H_2O$ |

Supplementary Table S3 details the various conditions that were examined in the development and defining the optimal liquid chromatography parameters. Initially, 2HG standards were prepared in 80% MeOH and dried under $N_2$ gas stream at room temperature. Dried standards were resuspended in 50 µL 200 mM chiral derivatization agent (CDA)—composed of DATAN, 4:1 ACN/Acetate and 2 µM sodium lactate) for 30 min at 70°C. Derivatized standards were dried under $N_2$ gas stream at 35°C. Standards were diluted in 50% ACN (aq). Initially, C18 columns (Xselect BEH phenyl C18, Xselect HSS T3, Eclipse Plus RRHD were compared over the reverse phase gradient composed of mobile phases: A (2 mM ammonium formate in $H_20$ +0.01%v/v 5 mM Medronate pH 3.5); B. MeOH. Reverse phase gradient was applied as described in Methods section. While two other polar stationary phase columns (Discovery HSF5 and Sequant HILIC-Z) were compared over a normal phase gradient composed mobile phases of A (20 mM FA pH 3.25 in $H_20$) and B (90:10 ACN/20 mM FA pH 3.25). Normal phase gradient was run as indicated here: 85% B initial; hold 30 min; ramp to 50% B by 33 min; ramp to 25% B by 37 min. Comparison with the AdvanceBio Glycan Map (an amide base HILIC column) was used based on alternated in-house normal phase gradient as described in Supplementary Figure S6 and S7. LC/MS ESI- mode parameters were applied as described in Methods section (unless otherwise indicated). Upon determining that Zorbax EclipsePlus RRHD provided improved separation and retention over Xselect HSS T3 C18, further steps were conducted to optimize chemistry to promote enantiomeric separation and enhanced signal detection of derivatized D-2HG/L-2HG analogs from 2HG-DATAN esters. During this phase of method optimization, qualitative comparison of the enantiomeric separation of 2HG following various changes in the compositions of buffers, additives and diluents w (detailed in Supplementary Table S2) were conducted using the Zorbax EclipsePlus RRHD column. In-depth details for comparisons that were not substantially different or leading to enhancements are not discussed or illustrated (aside from the representative HILIC comparisons as shown in Supplementary Figure S6 and S7, below) due to their low relevance to the objective of improving chromatography.

**Supplementary Table S4.** QTOF-MS/MS analysis with commercial standards for targeted TOF-MS reference library confirmation following chiral derivatization. *

| Analyte ID | Precursor | Ret. Time | CID transition |
|---|---|---|---|
| | m/z | min | m/z |
| D-2-Phosphoglycerate | 184.986 | 1.7 | 96.961, 166.977, 184.986 |
| D-3-Phosphoglycerate | 184.986 | 2.7 | 78.960, 96.961, 184.986 |
| L-Arginine | 173.104 | 1.2 | 131.081, 173.104 |
| L-Asparagine | 131.046 | 1.5 | 95.025, 113.035, 114.019, 131.041 |
| L-Aspartate | 132.030 | 1.4 | 72.012, 88.101, 115.001, 133.011 |
| Citrate | 191.019 | 1.3 | 101.024, 111.006, 117.017, 129.017, 191.019 |
| D-Glyceraldehyde 3-phosphate | 168.990 | 1.6 | 78.959, 96.970, 150.981, 168.991 |
| D-Diphosphoglyceraterate | 264.952 | 1.4 | 264.952, 246.943, 166.975 |
| D-Lactate | 89.024 | 7.5 | 71.013, 89.024 |
| D-Malate | 133.014 | 3.5 | 115.003, 133.013 |
| FAD, Flavin adenine dinucleotide | 784.150 | 10.5 | 346.055,437.089,784.149 |
| FMN, Flavin mononucleotide | 455.097 | 10 | 78.960, 96.970, 199.003, 213.018, 445.099 |
| D-Glucose | 179.056 | 1.4 | 71.014, 89.024, 101.025,179.0558 |
| L-Glutamate | 146.046 | 1.3 | 102.056, 128.036,146.048 |
| L-Glutamine | 145.062 | 1.25 | 127.051,128.037, 145.062 |
| GSSG, Oxidized glutathione disulfide | 611.145 | 3 | 143.046, 254.077, 272.088,306.077, 611.1455 |
| DL-β-Hydroxybutyrate | 103.040 | 9.8 | 59.014, 103.040 |
| Isocitrate | 191.019 | 1.9 | 85.026, 101.0235, 111.0061, 117.0171, 129.0169, 191.019 |
| L-Leucine | 130.087 | 3.1 | 130.088** |
| L-Lactate | 89.024 | 7 | 71.013, 89.024 |
| L-Malate | 133.014 | 3.1 | 115.003, 133.013 |
| L-Phenylalanine | 164.072 | 6 | 147.042, 164.071 |
| L-Lysine | 145.098 | 1.1 | 99.0918,145.098 |
| PEP, Phosphenolpyruvate | 166.975 | 1.4 | 78.9590,166.975 |
| Pyridoxine | 168.066 | 9.8 | 138.058, 150.057, 168.066 |
| Succinate | 117.019 | 2.7 | 73.026, 99.008, 117.019 |
| Taurine | 124.008 | 1.4 | 79.957,124.007 |
| L-Tyrosine | 180.066 | 2.8 | 93.036, 119.048, 163.038, 180.067 |
| L-Tryptophan | 203.082 | 9.9 | 116.05, 142.066, 159.091,203.084 |
| L-Isoleucine | 130.087 | 3.2 | 130.091** |

*Supplementary Table S4 provides MS/MS–based validation was applied to produce fragmentation fingerprints which were generated using the LC/MS conditions outlined in the Methods section and collision-induced dissociation (CID) by ramping the collision energy (CE) from 0 – 10 V. Commercial standards were purchased from Sigma and prepared following the derivatization and LC/MS conditions outlined in the Methods section. Each analyte identity is provided along with their product ion and transition m/z values can are provided in MS/MS–based validation All fragmentation patterns were verified transition against the online HMDB and Metlin MS/MS reference library. **Refers to m/z that did not have additional fragments in the online MS/MS reference libraries.

**Supplementary Table S5.** Statistic results Table for the comparative analysis of representative IDH1$^{wt}$ versus IDH1$^{mut}$ tumor spheroids comparison for cells grown under normoxic conditions.

| Name | Response | P-value | FC | log (FC) |
|---|---|---|---|---|
| D-2HG | ↓ IDH1$^{wt}$, ↑IDH1$^{mut}$ | 4.3E-08 | 0.01 | -6.7 |
| Hydroxybutyrate | ↑ IDH1$^{wt}$, ↓IDH1$^{mut}$ | 8.6E-04 | 2.33 | 1.2 |
| L-2HG | ↑ IDH1$^{wt}$, ↓IDH1$^{mut}$ | 1.5E-03 | 1.92 | 0.9 |
| GSSG, Glutathione Disulfide, Oxidized | ↓ IDH1$^{wt}$, ↑IDH1$^{mut}$ | 5.8E-03 | 0.31 | -1.7 |
| Glutamine | ↑ IDH1$^{wt}$, ↓IDH1$^{mut}$ | 1.7E-02 | 4.35 | 2.1 |
| Succinate | ↓ IDH1$^{wt}$, ↑IDH1$^{mut}$ | 2.0E-02 | 0.04 | -4.5 |
| L-Lactate | ↑ IDH1$^{wt}$, ↓IDH1$^{mut}$ | 2.7E-02 | 1.55 | 0.6 |
| Phenylalanine | ↑ IDH1$^{wt}$, ↓IDH1$^{mut}$ | 3.3E-02 | 11.8 | 3.6 |
| Arginine | ↑ IDH1$^{wt}$, ↓IDH1$^{mut}$ | 3.5E-02 | 5.72 | 2.5 |
| Isocitrate | ↑ IDH1$^{wt}$, ↓IDH1$^{mut}$ | 5.9E-02 | 1.98 | 1.0 |
| L-Malate | ↓ IDH1$^{wt}$, ↑IDH1$^{mut}$ | 7.2E-02 | 0.61 | -0.7 |
| D-3-Phosphoglycerate | ↓ IDH1$^{wt}$, ↑IDH1$^{mut}$ | 9.2E-02 | 0.10 | -3.3 |

**Supplementary Table S6.** Matrix factor for our LC/MS D-2HG and L-2HG quantitation method*.

| Analyte | Mean Peak Area | RSD (%) | Matrix Factor |
|---|---|---|---|
| **L-2HG added, nmol/mL (n=3)** | | | |
| **0** | **7.0E+04** | **4.8** | |
| **13** | **7.5E+05** | **3.6** | **1.2** |
| **D-2HG added, nmol/mL (n=3)** | | | |
| **0** | **8.5E+04** | **3.4** | |
| **13** | **1.0E+06** | **5** | **0.9** |

*In Supplementary Table S6, matrix factor was determined for an IDH1 mutant (TS603) and IDH1 wild-type (GSC923) cells spiked with D-2HG and L-2HG that were prepared, derivatized with DATAN and analyzed by LC/MS as described in the Methods section. Matrix factor was determined based on area of standard/ (area of spiked-area - area of non-spiked).
